# Regulation of FLIP(L) and TRAIL-R2 signalling by the SCF^Skp2^ Ubiquitin Ligase Complex

**DOI:** 10.1101/723718

**Authors:** JZ Roberts, C Holohan, T Sessler, J Fox, C. Higgins, G Espona-Fiedler, J Majkut, N Crawford, JS Riley, H Khawaja, LM Humphreys, J Ferris, E Evergren, P Moynagh, SS McDade, DB Longley

## Abstract

Depending on its expression levels, the long splice form of the pseudo-caspase FLIP (FLIP(L)) can act as an inhibitor (high expression) or activator (low expression) of apoptosis induction by the TRAIL-R2 death-inducing signalling complex (DISC); its expression levels are therefore tightly regulated. Here, we demonstrate that the Skp1-Cullin-1-F-box (SCF) Cullin-Ring E3 Ubiquitin Ligase complex containing Skp2 (SCF^Skp2^) regulates the stability of FLIP(L) (but not the short splice form FLIP(S)), and, unusually, this is mediated by direct binding of FLIP(L) to Cullin-1 rather than via Skp2. By fine mapping the interaction of FLIP(L) with Cullin-1 to the large subunit of its pseudo-caspase domain, we found that the interaction is significantly stronger with FLIP(L)’s DISC-processed p43-form. Importantly, this interaction disrupts the ability of p43-FLIP to interact with FADD, caspase-8 and another DISC component, TRAF2. Moreover, we find that SCF^Skp2^ associates with TRAIL-R2 constitutively and does so independently of FLIP(L) and other canonical DISC components. Inhibition of Cullin-1 expression (using siRNA) or activity (using a NEDDylation inhibitor, MLN4924) enhanced FLIP(L) and TRAF2 levels at the TRAIL-R2 DISC and enhanced caspase-8 processing. This suggests that processing of FLIP(L) to p43-FLIP at the TRAIL-R2 DISC enhances its interaction with co-localised SCF^Skp2^, leading to disruption of p43-FLIP’s association with the DISC thereby altering caspase-8 processing. These findings provide important new insights into how FLIP(L) expression and TRAIL-R2 signaling is controlled.

## Introduction

Programmed cell death (apoptosis) plays a key role in maintaining normal tissue homeostasis and preventing disease^1^. Apoptosis is orchestrated by a family of cysteine proteases, the caspases^2^. The multi-protein complexes formed following activation of the CD95 (Fas) and TRAIL-R1/R2 (DR4/DR5) death receptors by their ligands (FasL and TRAIL) expressed by immune effector cells and by 2^nd^ generation therapeutic agonists (ABBV621 and MEDI3039) is called the death-inducing signalling complex (DISC), consisting of the receptors, the adaptor molecule FADD, procaspase-8 and FLIP^3^. Recruitment into these complexes exposes FADD’s N-terminal death effector domain (DED), which recruits procaspase-8 by interacting with its N-terminal tandem DEDs^4^. Procaspase-8 homodimerization results in conformational changes in its catalytic domains that lead to its activation^5^. Two main splice forms of FLIP have been identified in humans: a long form (FLIP(L)) and a short form (FLIP(S)), both of which contain tandem DEDs and can be recruited to the DISC and related complexes (such as TNFR1 Complex II and the ripoptosome) where they form heterodimers with procaspase-8^4^. The relative amounts of FLIP(S) and FLIP(L) recruited to these complexes is a key determinant of cell fate (survival, apoptosis or necroptosis) and whether FLIP(L) acts as an inhibitor or activator of caspase-8-mediated apoptosis in these complexes depends on its stoichiometry relative to caspase-8; therefore FLIP(L) expression is tightly controlled at multiple levels^3,4^.

It has been established that FLIP is regulated through the Ubiquitin Proteasome System (UPS), although the regulation of the two major splice forms of FLIP is distinct. The UPS degrades FLIP(S) more rapidly than FLIP(L) due to a unique C-terminal tail possessed by FLIP(S). Lys^192^ and Lys^195^ are the main ubiquitination sites on FLIP(S)^6^, and it was later found by the same group that phosphorylation of Ser^193^ decreases FLIP(S) ubiquitination and increases its half-life^7^. It has also been reported that the PTEN/Akt pathway can regulate FLIP(S) ubiquitination through an ubiquitin E3 ligase named ITCH (AIP4)^8^, which has also been reported to directly ubiquitinate FLIP(L)^9,10^. Hyperthermic stress^11^ or ROS induction^12^ have both been shown to down-regulate FLIP(L) at the protein level; mutation of Lys^195^ blocked hyperthermia-induced down-regulation, whereas mutation of Lys^167^ and an adjacent phosphorylation site (Thr166) inhibited its ROS-mediated down regulation.

Other ubiquitin-like proteins (UBL) are conjugated to substrates in a similar manner to ubiquitin. NEDD8 is a 9kDa UBL first discovered in 1992, when it was shown to be downregulated during mouse brain development^13,14^. Just like the ubiquitination cascade, the NEDDylation cascade is an ATP-dependent process that relies on an E1/E2/E3 enzyme system; this results in the formation of an isopeptide bond between the C-terminus of NEDD8 (Gly76) and a substrate’s lysine residue. To date, relatively few proteins have been identified to be NEDDylated, with the most researched NEDDylation substrates being the Cullin-Ring E3 Ligase (CRL) family of E3 ubiquitin ligases^15,16^. The CRLs are multimeric ubiquitin E3 ligases made up of three key sections (**Figure 1A**): the Cullin, which normally acts as a scaffold between the C-terminal catalytic region and the N-terminal substrate recognition region of the CRL; the RING finger domain-containing Rbx1/2 catalytic subunit; and the substrate-binding domain, which consists of an adaptor (that binds the Cullin) and substrate receptor (that binds the target protein). The activity of CRLs is critically dependent on NEDDylation: when a CRL is NEDDylated, it is active.

**Figure 1.**
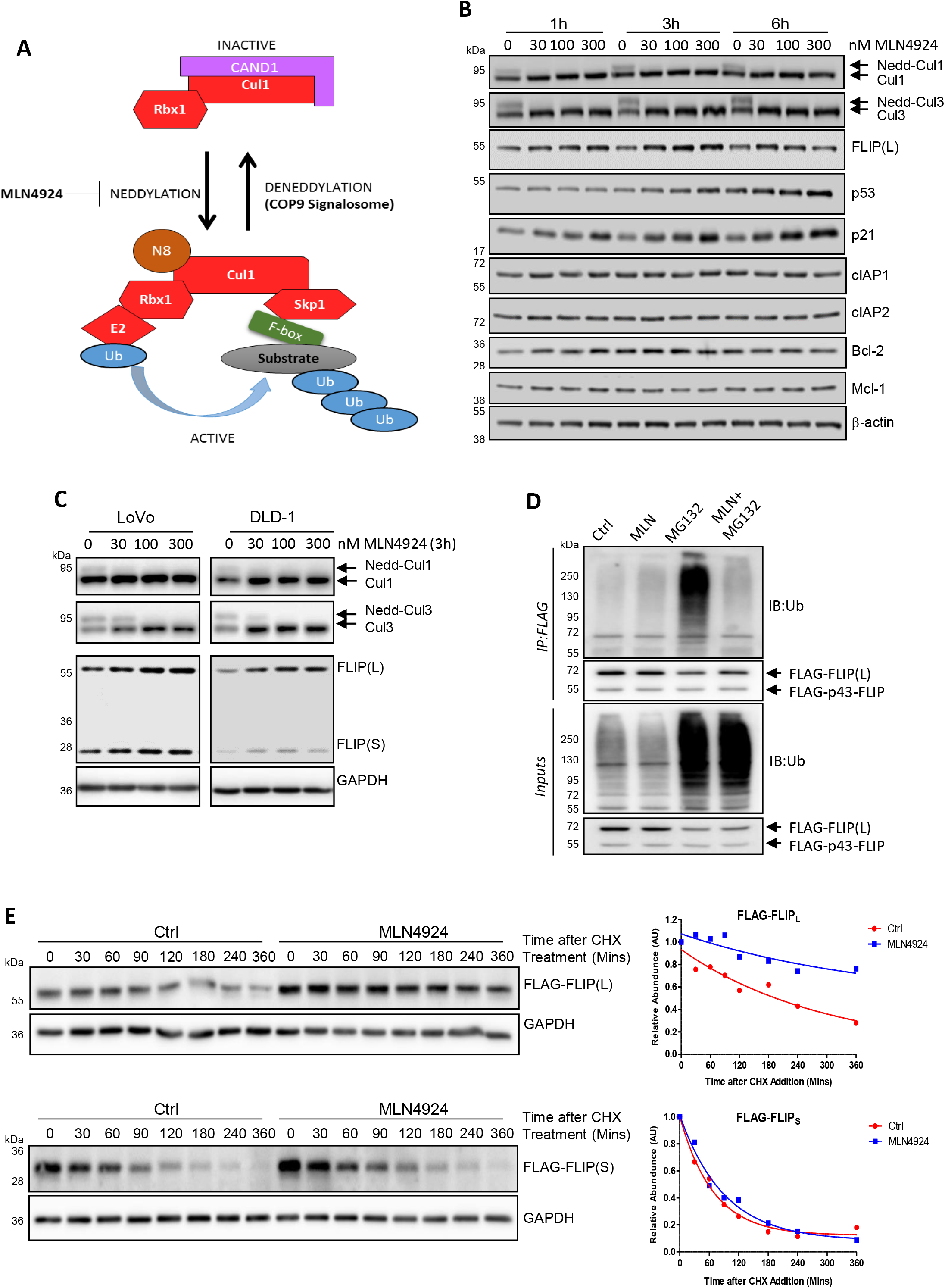
FLIP(L) ubiquitination and protein stability is NEDDylation-dependent. **(A)** Schematic diagram of the Skp1-Cullin-1-F-box protein (SCF) Cullin-RING Ligase (CRL) complex and its regulation by NEDDylation. **(B)** Western blot analysis of apoptotic protein expression and Cullin-1 and −3 expression in HCT116 cells treated with MLN4924. **(C)** Western blot analysis of FLIP and Cullin-1 and −3 expression in LoVo and DLD-1 cells treated with MLN4924. **(D)** FLAG-tagged FLIP(L) was transiently overexpressed in HEK293T cells, which were then treated with MLN4924 (300nM) for 1h before treatment with the proteasome inhibitor MG132 (10μM) for an additional 2h. FLAG-FLIP(L) was immunoprecipitated under denaturing conditions and a Western blot performed for ubiquitin (Ub). **(E)** Protein half-life experiments conducted in HCT116 cells stably expressing either FLAG-tagged FLIP(L) or FLAG-tagged FLIP(S) treated with 100ug/mL of the protein synthesis inhibitor cycloheximide (CHX).

In this study, we identify the Skp1-Cullin-1-F-box protein containing Skp2 complex (SCF^Skp2^) as a specific regulator of ubiquitination and stability of the long FLIP splice form that also interacts with TRAIL-R2. Moreover, we find that FLIP(L) directly interacts with Cullin-1 rather than via Skp2 and that this interaction is significantly enhanced when FLIP(L) is in its DISC-processed p43-form. Not only does this interaction regulate FLIP(L) turnover, it also disrupts the interaction of FLIP(L) with FADD and caspase-8 and decreases its levels at the TRAIL-R2 DISC. These results identify a unique mechanism by which expression of FLIP(L) and its interaction with and regulation of the TRAIL-R2 DISC is controlled.

## Results

### FLIP(L) ubiquitination and protein stability is NEDDylation-dependent

MLN4924 (TAK4924/ Pevonedistat) is a selective, small molecule inhibitor of the NEDD8-activating E1 enzyme (NAE1) under clinical evaluation for the treatment of a number of cancers^17^. In this study, we exploited the knowledge that inhibition of NAE1 blocks Cullin NEDDylation, thereby rendering CRLs inactive (**Figure 1A**). Rapid loss of NEDDylated Cullin-1 and Cullin-3 was observed in response to 30nM MLN4924, and the known CRL substrate, p21, increased in a concentration-dependent manner (**Figure 1B**). Of the apoptotic proteins profiled, only FLIP(L) was found to be acutely upregulated in response to MLN4924 (**Figure 1B; Supplementary Figure 1A** *and data not shown*); FLIP(S) was below the level of detection in these studies. This upregulation of FLIP(L) was also observed at 300nM MLN4924 at 3h, although this was lost by 6h. FLIP mRNA levels were not affected by MLN4924 treatment (**Supplementary Figure 1B**), indicating that the observed increase in FLIP(L) levels was post-transcriptional. Similar effects were observed in the p53 wild-type LoVo and p53 mutant DLD-1 CRC models (**Figure 1C** and **Supplementary Figure 1C-E**), indicating that the effects of MLN4924 on FLIP(L) are post-transcriptional and not p53-dependent. Importantly, MLN4924 inhibited the accumulation of ubiquitinated FLIP(L) in cells co-treated with the proteasome inhibitor MG132 (**Figure 1D**), while in half-life studies, MLN4924 enhanced the stability of FLIP(L), but not FLIP(S) (**Figure 1E**). Collectively, these results indicate that FLIP(L)’s ubiquitination and turnover by the proteasome is regulated by MLN4924.

### FLIP(L) ubiquitination is regulated by the Skp1-Cullin-1-F-box protein containing Skp2 (SCF^Skp2^) complex

Previous studies have linked resistance to TRAIL-induced apoptosis to Skp2^18^, an F-box protein which is part of the substrate receptor for the SCF CRL complex (**Figure 1A**). Given its key role in regulating TRAIL-induced apoptosis, we assessed whether NEDDylation-dependent FLIP(L) ubiquitination was regulated by this complex. Notably, RNAi-mediated downregulation of Skp1, Cullin-1 or another key component of the SCF^Skp2^ complex, Rbx1, decreased basal FLIP ubiquitination (**Figure 2A**). We next used a model stably over-expressing FLIP(L) to uncouple protein expression from regulation of transcription of the endogenous gene (FLIP is transcriptionally regulated by NFκB which is itself regulated by Cullin-1); we found that downregulation of Skp1, Cullin-1 or (to a lesser extent) Rbx1 enhanced FLIP(L) protein expression (**Figure 2B**). Similarly, RNAi-mediated downregulation of Skp2 enhanced endogenous FLIP(L) but not FLIP(S) protein expression, and this was most apparent when cells were co-treated with MG132 (**Figure 2C**). Additionally, Skp2 downregulation was found to inhibit endogenous FLIP ubiquitination (**Figure 2D**). Interestingly, FLIP(S) levels increased with siRNA-mediated depletion of Rbx1 (**Figure 2A**), the catalytic subunit shared with many of the CRL complexes, suggesting that another CRL complex might regulate FLIP(S) levels, potentially explaining why we observed an increase in FLIP(S) in response to MLN4924 treatment in the LoVo and RKO models (**Figure 1C**). Most importantly, we subsequently demonstrated that both Cullin-1 (**Figure 2E**) and Skp2 (**Figure 2F**) can interact with FLIP(L) but not FLIP(S) and that simultaneous over-expression of both Cullin-1 and Skp2 enhanced ubiquitination of endogenous FLIP, which was reversed with MLN4924 treatment (**Figure 2G**). Collectively, these results indicate that the SCF^Skp2^ complex can interact selectively with the long FLIP splice form and induce its ubiquitination.

**Figure 2.**
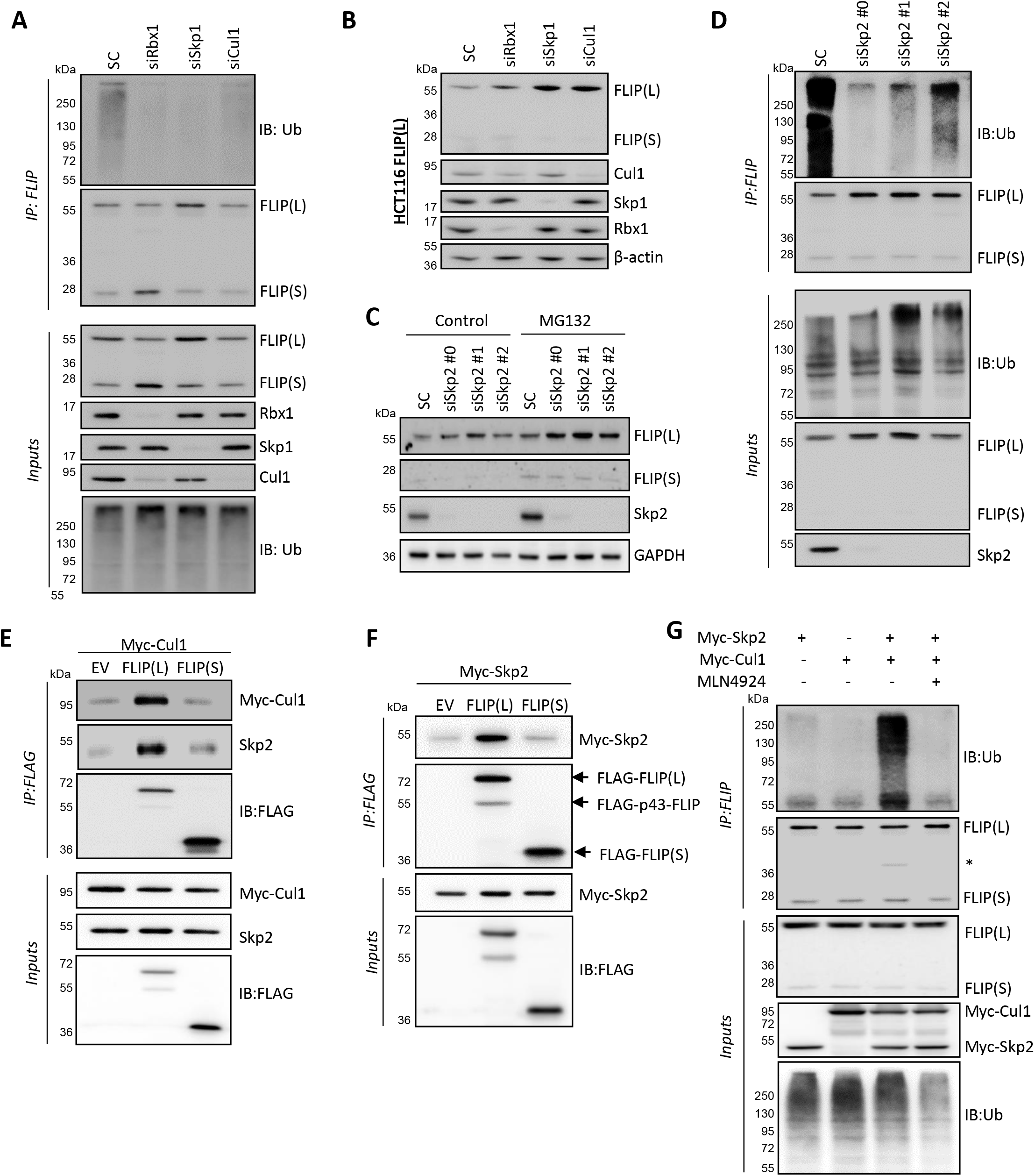
FLIP(L) ubiquitination is regulated by SCF^Skp2^. **(A)** Western blot of FLIP ubiquitination in HCT116 cells transfected for 48h with siRNAs (20nM) targeting Rbx1, Skp1 or Cullin-1. **(B)** Western blot analysis of exogenous FLIP(L) expression in HCT116 cells transfected for 48h with siRNAs (20nM) targeting Rbx1, Skp1 or Cullin-1. **(C)** Western blot analysis of FLIP(L) expression in HCT116 cells transfected for 48h with siRNAs (10nM) targeting Skp2; cells were treated for 2h with 10μM MG132 prior to collection as indicated. **(D)** Western blot analysis of FLIP ubiquitination in HCT116 cells transfected for 48h with siRNAs (10nM) targeting Skp2. **(E)** Co-immunoprecipitation (Co-IP) of Myc-tagged Cullin-1 with FLAG-tagged FLIP(L) or FLIP(S); interaction with endogenous Skp2 was also assessed. **(F)** Co-IP of Myc-tagged Skp2 with FLAG-tagged FLIP(L) or FLIP(S). **(G)** Impact of Cullin-1 and Skp2 overexpression on FLIP ubiquitination in the presence/absence of MLN4924 (300nM, 3h).

### Mapping the SCF^Skp2^ protein interaction site in FLIP(L)

Initial mapping studies localised the site of interaction between FLIP(L) and Cullin-1/Skp2 to a region between amino acids 211 and 350, which corresponds primarily to the large p20 subunit of its pseudo-caspase domain (**Supplementary Figure 2A-C**). This is consistent with lack of interaction with FLIP(S), which does not contain this domain. Further mapping narrowed the interaction site to a region between amino acids 255 and 292 (**Figure 3A-C**). In these experiments, it was noticeable that the interaction between FLIP(L) and Cullin-1/Skp2 was significantly enhanced by deletion of the C-terminal region of FLIP(L), which contains the p12-pseudo-caspase subunit that is cleaved in a caspase-8-dependent manner at the DISC (**Figure 3A-C** and **Supplementary Figure 2A-C**). We were subsequently able to fine map the Cullin-1/Skp2 interaction domain to a 13 amino acid region in FLIP(L) (**Figure 3D-F**) corresponding to amino acids 255-267 (**Figure 3G**). Skp2 normally binds to “phospho-degron” motifs on its substrates, which typically contain at least one phosphorylated Threonine or Serine^19^; however, phosphorylation of Thr264 was not required for FLIP(L)’s interaction with Cullin-1/Skp2 (demonstrated by mutation of this residue to Alanine; **Supplementary Figure 3A-D**). We also assessed whether Asp258, Glu263 or Glu265 were potentially acting as phospho-mimetic residues; however, mutation of these residues to alanine also failed to abrogate FLIP(L)’s interaction with Cullin-1/Skp2 (**Supplementary Figure 3E**). Nonetheless, we were able to show that the identified region of FLIP(L) was able to interact with all SCF^Skp2^ components, utilising a Biotin-tagged peptide incorporating the Cullin-1/Skp2 interaction region of FLIP(L) (plus additional flanking residues) (**Figure 3H**). Furthermore, using a model of the large and small catalytic subunits of the caspase-8/FLIP(L) heterodimer derived from its crystal structure^20^, we found that most of the 13 amino acid Cullin-1 interaction region (red) in the p20 subunit of FLIP(L) is present on the protein surface and therefore available for forming protein-protein interactions (**Figure 3I**). In addition, this region is adjacent to and interacts with FLIP(L)’s p12 subunit of its pseudo-catalytic domain (**Supplementary Figure 4C**). FLIP(L) is cleaved between its large (p20) and small catalytic subunits by caspase-8 when it forms a heterodimer with caspase-8 at the DISC and related complexes; however, it is important to note that the small subunit remains associated with the heterodimer and is essential for its catalytic activity^20^.

**Figure 3.**
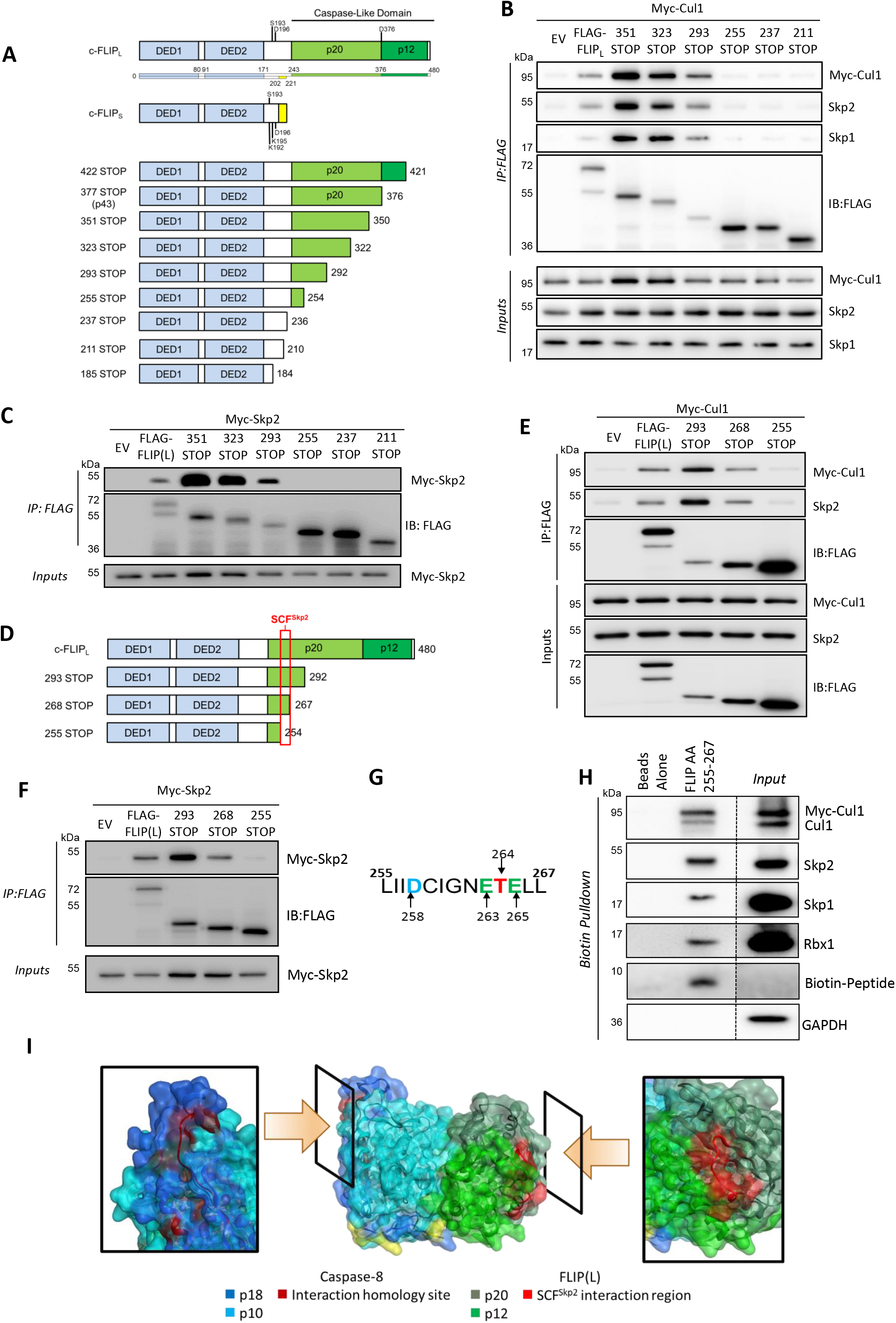
Mapping SCF^Skp2^ interaction site within FLIP(L). **(A)** Schematic diagram of FLIP expression constructs; the death effector domains (DEDs) and large (p20) and small (p12) subunits of the pseudo-caspase domain are highlighted. Co-IP experiment mapping the interaction site between FLIP(L) and **(B)** Cullin-1 and **(C)** Skp2. **(D)** Schematic diagram of FLIP expression constructs; the region of interaction with the SCF^Skp2^ complex is highlighted. Co-IP experiment fine mapping the interaction site between FLIP(L) and **(E)** Cullin-1 and **(F)** Skp2. **(G)** Amino acid sequence of FLIP(L)/SCF^Skp2^ complex interaction site, and **(H)** biotinylated peptide pull-down experiment demonstrating the interaction of this region with the SCF^Skp2^ complex. **(I)** Model derived from the crystal structure (PDB ID: 3H11) of the large and small catalytic domains of the caspase-8/FLIP(L) heterodimer. The position of the SCF^Skp2^ complex interaction site and the equivalent (non-contiguous) residues in caspase-8 are highlighted in red. A frontal view of the heterodimer interface is depicted in the middle; the “windows” show 90<u>°</u> rotations of the crystal structure, highlighting the SCF^Skp2^ interaction of FLIP(L) and its equivalent region in Procaspase-8.

### FLIP(L) cleavage promotes its interaction with the SCF^Skp2^ complex

Given the proximity of FLIP(L)’s SCF^Skp2^ interaction site to its p12-domain (**Figure 3I** and **Supplementary Figure 4C**), we investigated the effect of FLIP(L)’s cleavage for its interaction with the SCF^Skp2^ complex. A C-terminal deletion corresponding to p43-FLIP (the caspase-8 cleavage product typically detected at the DISC) and a non-cleavable form of FLIP(L) (NC-FLIP(L), D376A) were generated. Notably, p43-FLIP interacted more strongly with Cullin-1, Skp1 and Skp2 than the wild-type protein, whereas the NC-FLIP(L) interacted to the same extent (**Figure 4A**). In addition, NC-FLIP(L) was significantly more stable than FLIP(L) and p43-FLIP (**Figure 4B**). These results suggest that cleavage of FLIP(L) to its p43-form promotes its interaction with the SCF^Skp2^ complex (potentially by relaxing the p20/p12 interface) and subsequent degradation via the UPS. As FLIP(L) cleavage to its p43-form is mediated by caspase-8 in a FADD-dependent manner, we assessed whether these canonical DISC components could also interact with the SCF^Skp2^ complex. However, neither procaspase-8 nor FADD (nor as expected FLIP(S)) co-immunoprecipitated with Cullin-1 (**Figure 4C**). This is consistent with the fact that the 13 amino acid region in FLIP(L) that interacts with Cullin-1 is not conserved in procaspase-8, which has a 21 amino acid insertion in this region (**Figure 3I** and **Supplementary Figure 4A-B**).

**Figure 4.**
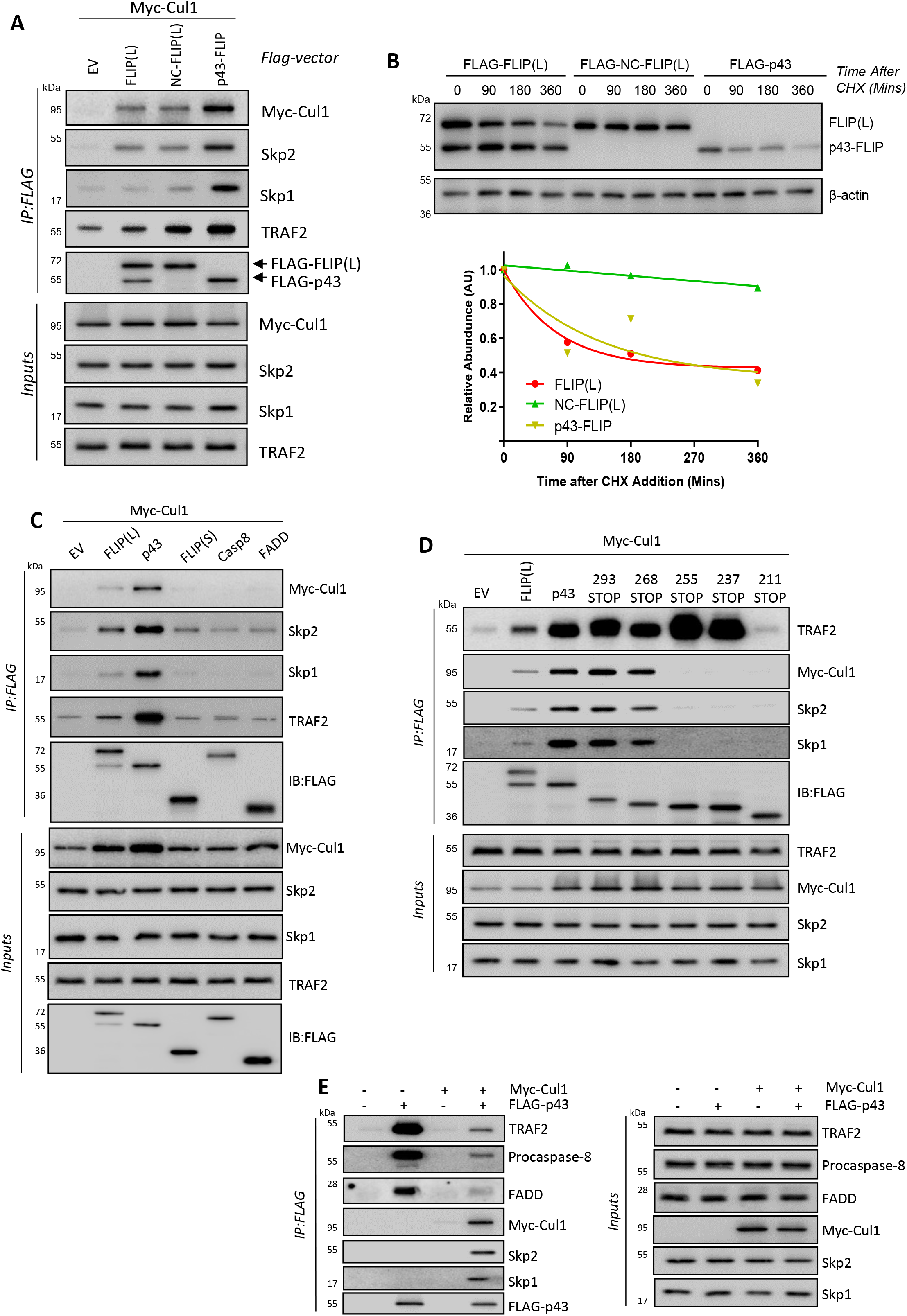
FLIP(L) cleavage promotes its interaction with SCF^Skp2^. **(A)** Co-IP analysis of the interaction of Cullin-1 with wild-type, non-cleavable (NC)-FLIP(L) and p43-FLIP. Interactions with endogenous Skp1, Skp2 and TRAF2 were also assessed. **(B)** Western blot assessment of half-lives of wild-type-, NC- and p43-FLIP(L) using 100ug/mL cycloheximide (CHX). **(C)** Co-IP analysis of the interaction of Cullin-1 with full-length FLIP(L), p43-FLIP(L), FLIP(S), caspase-8 (Casp8) and FADD; interactions with endogenous Skp1, Skp2 and TRAF2 were also assessed. **(D)** Co-IP analysis of the interaction of Cullin-1 with full-length and truncated forms of FLIP(L); interactions with endogenous Skp1, Skp2 and TRAF2 were also assessed. **(E)** Co-IP analysis of the interaction of p43-FLIP with endogenous FADD, caspase-8 and TRAF2 in the presence and absence of overexpressed Cullin-1; endogenous Skp1 and Skp2 were also assessed.

TRAF2 is another DISC-recruited protein that also plays a key role in death receptor signalling^21,22^. Moreover, TRAF2 has been reported to interact with FLIP(L) and (more strongly) p43-FLIP^23^, which we confirmed (**Figure 4A**). Mapping studies re-confirmed the location of the Cullin-1:FLIP(L) interaction site and demonstrated that the TRAF2:FLIP(L) interaction site is located in an adjacent region, between amino acids 211 and 236 (**Figure 4D** and **Supplementary Figure 4D**). TRAF2 co-immunoprecipitated with Cullin-1 in the presence of p43-FLIP or FLIP(L) and did not interact with caspase-8 or FADD (**Figure 4C**), even though TRAF2 has been reported to act as an E3 ligase for caspase-8 at the DISC^24^. Moreover, in co-transfection experiments, the interaction of p43-FLIP with TRAF2, FADD and procaspase-8 was significantly reduced in cells co-transfected with Cullin-1 (**Figure 4E**), suggesting competition between these canonical DISC proteins and Cullin-1 for binding to p43-FLIP. In complementary co-transfection experiments, procaspase-8 disrupted the interaction of p43-FLIP with Cullin-1 (**Supplementary Figure 5A/B**).

### Mapping the FLIP(L) interaction site in the SCF^Skp2^ complex

The above results (**Figure 3**) indicate that FLIP(L) does not interact with the SCF^Skp2^ complex in a canonical manner; *i.e*. via a phospho-degron motif. We therefore assessed the requirement of Skp1 and Skp2 for FLIP(L)’s interaction with Cullin-1. In these experiments, we focussed on the more strongly interacting p43-FLIP. Notably, RNAi-mediated down-regulation of either Skp1 or Skp2 failed to abrogate the p43-FLIP:Cullin-1 interaction, although Skp1 silencing disrupted the interaction of Skp2 with p43-FLIP and Cullin-1 (**Figure 5A**), suggesting that Skp2’s interaction with FLIP(L) is indirect and mediated by Cullin-1 via Skp1. Skp2 normally interacts with substrates via its leucine-rich repeat (LRR) domain and with Skp1 via its F-box domain; however, deletion of the LRR domain failed to disrupt p43-FLIP’s interaction with Skp2, and the same was demonstrated with endogenous FLIP(L) (**Figure 5B**), further suggesting that the interaction between FLIP(L) and Skp2 is indirect.

**Figure 5.**
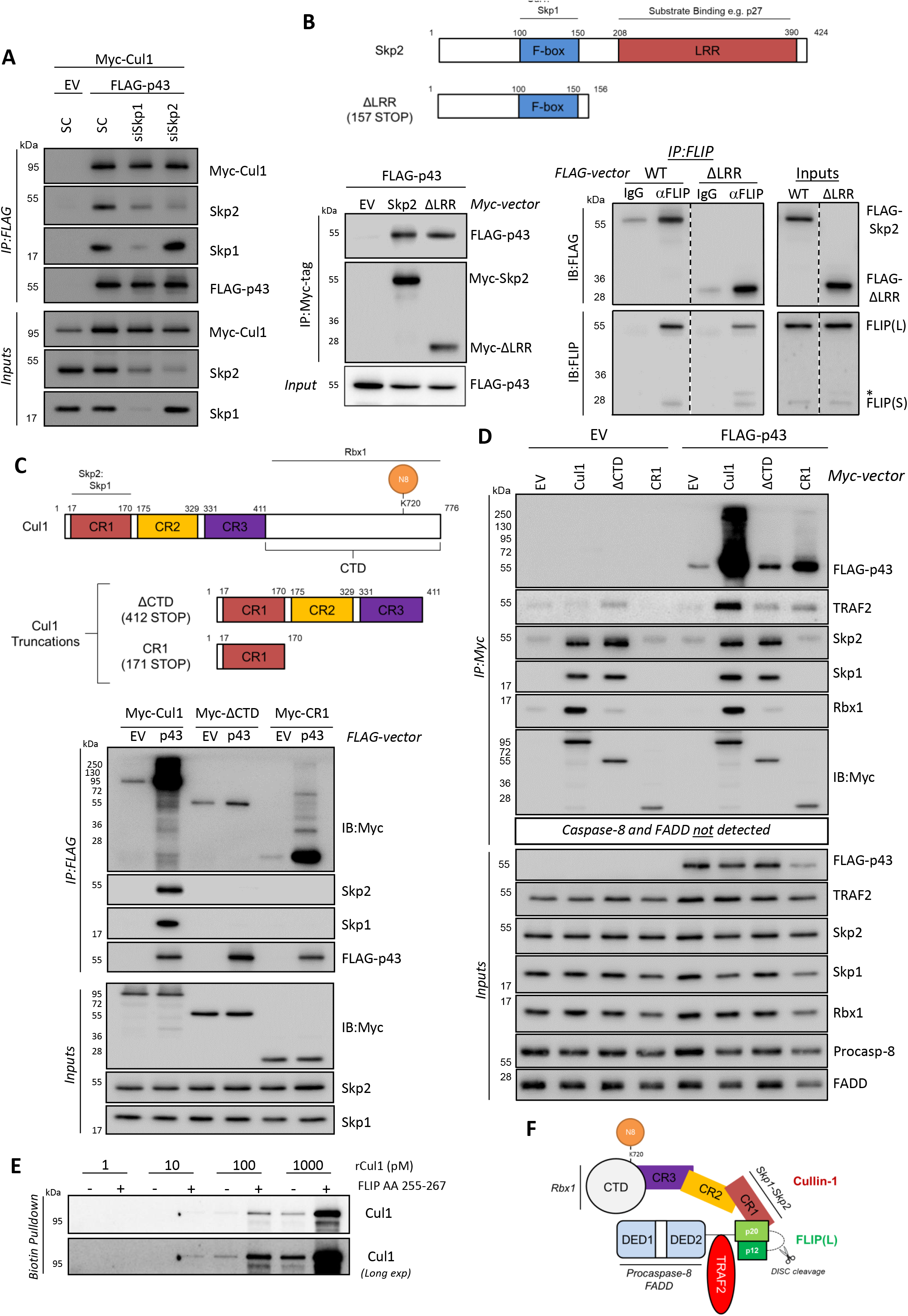
Mapping the interaction site of FLIP(L) within SCF^Skp2^. **(A)** Co-IP analysis of the interaction of p43-FLIP with Cullin-1, Skp1 and Skp2 in cells co-transfected with 10nM control (SC) or siRNAs targeting Skp1 or Skp2 for 48h. **(B)** Schematic diagram of full-length Skp2 and a C-terminal truncation that lacks its leucine-rich repeat (LRR) region. (*left panel*) Co-IP analysis of the interaction of p43-FLIP with full-length and truncated Skp2. (*right panel*) IP of endogenous FLIP showing its interaction with both full-length and truncated Skp2. **(C)** Schematic diagram of full-length Cullin-1 and two C-terminal truncations. Co-IP analysis of the interaction of p43-FLIP with full-length and truncated Cullin-1 and endogenous Skp1 and Skp2. **(D)** Co-IP analysis of the interaction of p43-FLIP with full-length and truncated Cullin-1 and endogenous Rbx1, Skp1 and Skp2. Components of the DISC were also assessed; neither caspase-8 nor FADD were detected, however TRAF2 was only detectable in cells co-transfected with p43-FLIP. **(E)** Interaction of recombinant Cullin-1 (rCul1) with biotin-linked peptide of FLIP(L) amino acids 255-267. **(F)** FLIP(L)’s interaction sites with TRAF2 and Cullin-1.

We next generated two C-terminal deletions of Cullin-1 both lacking its C-terminal domain (CTD); one containing all 3 Cullin-Repeat (CR1-3) domains (ΔCTD), and one containing only the N-terminal CR1 domain (**Figure 5C**). We found that the CR1 domain interacted with p43-FLIP (**Figure 5C-D**); however, although this has been reported to be the Skp1-interacting domain^25^, we found that the CR1 domain did not interact with Skp1 (or Skp2) in isolation from CR2/3 (**Figure 5D**), providing further evidence that FLIP(L) does not interact with Cullin-1 via Skp1/Skp2. Surprisingly, the interaction between p43-FLIP and the CR1-containing ΔCTD truncation was significantly weaker than the CR1-only truncation (**Figure 5C-D**); the reason for this is unclear, but it may be that when the globular Cullin-1 CTD is deleted, non-physiological interactions between the three CR domains disrupt the ability of the CR1 domain to interact with p43-FLIP. It is also worth noting that the fraction of p43-FLIP associated with full-length Cullin-1 (and therefore Skp1, Skp2 and Rbx1) was extensively modified, potentially by polyubiquitin chains (**Figure 5D**). Moreover, neither caspase-8 nor FADD were detected in this experiment, further confirming that these proteins do not interact with Cullin-1; however, an interaction with TRAF2 was detected in the presence of p43-FLIP. Importantly, we were able to confirm a direct interaction between Cullin-1 and FLIP(L) by incubating recombinant Cullin-1 (rCul1) with the Biotin-tagged peptide corresponding to the mapped interaction site on FLIP(L) (amino acid residues 255-267; **Figure 5E**). Collectively, the above results indicate that FLIP(L) directly interacts with the CR1 domain of Cullin-1 and that, unlike FADD and caspase-8, TRAF2 can potentially interact indirectly with Cullin-1 via FLIP(L) (**Figure 5F**).

### NEDDylated Cullin-1 is associated with the TRAIL-R2 DISC

Using a “DISC IP” assay for stimulating TRAIL-R2 DISC formation and then assessing the components of the complex^26^, we could detect Cullin-1, Rbx1, Skp1 and Skp2 associated with the TRAIL-R2 DISC in HCT116 p53 wild-type cells (**Figure 6A**). Notably, the levels of the SCF^Skp2^ complex recruited to the TRAIL-R2 DISC were higher in p53-deficient HCT116 cells, an effect not attributable to higher levels of expression of the complex’s components (**Supplementary Figure 6A**), suggesting a role for p53 in complex formation. Very low, if any, SCF^Skp2^ complex components could be detected in the DISCs formed by LIM1215 and COLO205 cells (**Figure 6A**), although these models expressed low levels of the SCF^Skp2^ complex (**Supplementary Figure 6A**) and cell surface TRAIL-R2 (as indicated by the amount of TRAIL-R2 immunoprecipitated). The levels of interaction between SCF^Skp2^ and TRAIL-R2 DISC in the HT29 model were similar to those in the HCT116 models; however, although the RKO model expressed all the SCF^Skp2^ components to a similar level as HCT116 and HT29 cell lines (**Supplementary Figure 6A**), very little SCF^Skp2^ was detected at the DISC despite significant cell surface TRAIL-R2 expression (**Figure 6A**). The levels of caspase-8 and FLIP at the DISC were also low at the RKO TRAIL-R2 DISC despite similar levels of expression (**Supplementary Figure 6A**) and similar levels of FADD recruitment (**Figure 6A**) to the other models. Of note, we also detected another CRL family member, Cullin-3, in these experiments; this is consistent with previous findings which reported that Cullin-3 promotes ubiquitin-dependent, DISC-mediated caspase-8 activation^27^. Importantly, despite similar expression of all SCF^Skp2^ components (**Supplementary Figure 6B**), none of the components of the SCF^Skp2^ complex were detected in control experiments conducted in cells in which TRAIL-R2 had been deleted (**Figure 6B**), confirming the specificity of the interaction. However, in cells lacking FADD, in which neither caspase-8 nor FLIP were recruited, Cullin-1 and the rest of the complex were detected at the DISC (**Figure 6B**), suggesting that the SCF^Skp2^ complex can interact with TRAIL-R2 in a FLIP-independent manner. This was further suggested by timecourse studies, in which components of the SCF^Skp2^ complex (and TRAF2) were all found to be associated with the unstimulated TRAIL-R2 (**Figure 6C**; 0 min). Moreover, stimulation of TRAIL-R2 caused an initial loss of the receptor’s association with SCF^Skp2^, followed by re-association with similar dynamics as TRAF2 and canonical DISC proteins, FLIP, FADD and caspase-8.

**Figure 6.**
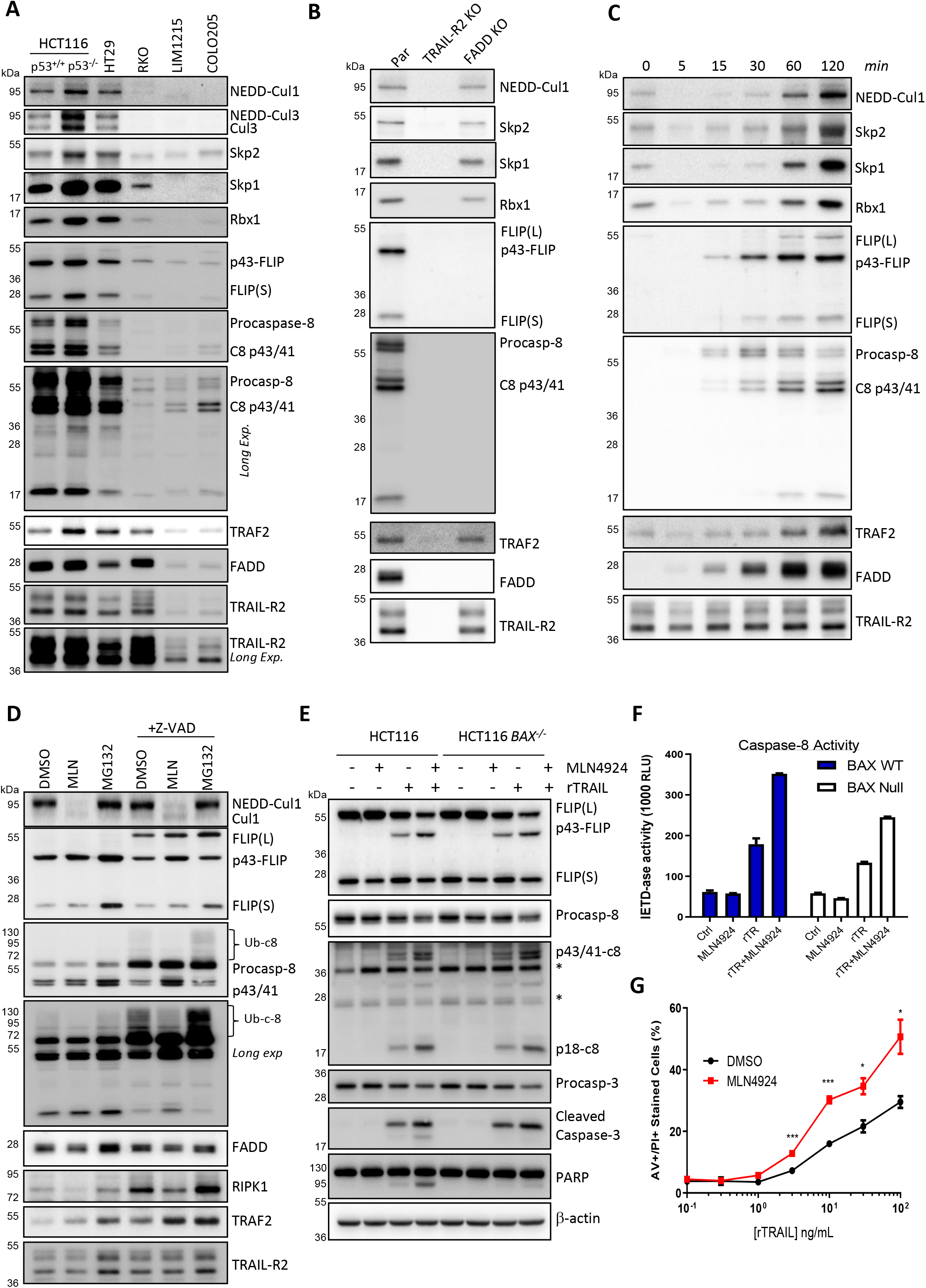
NEDDylated Cullin-1 is associated with the TRAIL-R2 DISC. **(A)** Western blot analysis of TRAIL-R2 DISC IP (1h) in a panel of colorectal cancer cell lines. **(B)** Western blot analysis of TRAIL-R2 DISC IP (1h) in parental and TRAIL-R2 and FADD CRISPR knockout HCT116 cell lines. **(C)** Timecourse Western blot analysis of TRAIL-R2 DISC IP in HCT116 cells. **(D)** Western blot analysis of TRAIL-R2 DISC IP in HCT116 cells treated for 1h with MLN4924 (100nM) or MG132 (10μM) prior to activation of the TRAIL-R2 receptor for 1h; cells were pre-treated with 20μM z-VAD-fmk pan-caspase inhibitor for 2h as indicated. Western blot analyses **(E)** and **(F)** caspase-8 activity assays in HCT116 parental and Bax-deficient cells co-treated with MLN4924 (100nM) and rTRAIL (10ng/mL) for 6h. **(G)** High content microscopy analysis of Annexin V/PI staining in HCT116 cells co-treated with MLN4924 (100nM) and rTRAIL (10ng/mL) for 24h. Significance was calculated by a Two-way ANOVA with a Bonferroni post hoc test.

Treating HCT116 cells with MLN4924 prior to DISC stimulation demonstrated that the form of Cullin-1 recruited to the DISC is its NEDDylated active form (**Figure 6D**). Furthermore, NEDDylated Cullin-1 was detected in membrane fractions, which were also enriched in TRAIL-R2 and canonical DISC and SCF^Skp2^ components (**Supplementary Figure 5C**). Notably, FLIP(L) was lost from the membrane fraction and enriched in the Lipid Raft/Cytoskeletal fraction following MLN4924 treatment (**Supplementary Figure 5C**); this was not observed for the other DISC proteins, further supporting FLIP(L)-specific regulation by the SCF^Skp2^ complex. Interestingly, nuclear levels of FLIP(L) were also enhanced in response to MLN4924 treatment (**Supplementary Figure 5C**); we and others have detected FLIP(L) in the nucleus^28–30^, where it has been reported to modulate important signalling pathways, for example Wnt^31^ and AP-1^32^.

Proteasome inhibitors have previously been reported to enhance DISC-mediated caspase-8 activation by inhibiting TRAF2-mediated ubiquitination of its large catalytic subunit^24^. Given their roles in inhibiting UPS-mediated protein degradation at different points, we compared the impact of MLN4924 and MG132 on TRAIL-R2 DISC assembly (**Figure 6D**). Treatment with both MLN4924 and MG132 enhanced the levels of both p43-FLIP and TRAF2 detected at the TRAIL-R2 DISC (~1.7-fold and ~1.9-fold respectively when quantified relative to TRAIL-R2). MG132 but not MLN4924 also enhanced FLIP(S) levels at the DISC. Another notable effect of MLN4924 was to reduce the levels of poly-ubiquitinated caspase-8 detectable at the DISC; this is consistent with inhibition of Cullin-3-mediated caspase-8 ubiquitination as previously reported^24^. Treatment with MLN4924 also decreased the levels of RIPK1 detected at the DISC, a substrate of the DISC-bound FLIP(L):caspase-8 heterodimer as well as caspase-8 homodimers. In the presence of the pan-caspase inhibitor zVAD-fmk, it was easier to visualise the effects of MLN4924, with enhanced levels of p43-FLIP and TRAF2, decreased levels of poly-ubiquitinated caspase-8 and decreased levels of RIPK1 again observed. Due to its ability to partially inhibit DISC-bound caspase-8, unprocessed FLIP(L) was detected in the presence of zVAD-fmk, and both MLN4924 and MG132 enhanced the levels of this form of FLIP(L) present in the complex.

To determine the net effect of these MLN4924-induced changes in DISC composition on TRAIL-induced apoptosis, we co-treated HCT116 cells with TRAIL and MLN4924 for 6h; enhanced levels of the active subunit of caspase-8 (p18) were detected by Western blot, and this correlated with enhanced PARP cleavage and caspase-8 activity (**Figure 6E/F**); similar results were also observed in cells pre-treated with MLN4924 (**Supplementary Figure 7A**). In addition, the levels of p43-FLIP and p43/41-caspase-8 were elevated in MLN4924/TRAIL co-treated cells (**Figure 6E**). Moreover, the effects of MLN4924 on caspase-8 and FLIP(L) processing were not due to downstream apoptosis signalling events as almost identical effects were observed in BAX-deficient cells (**Figure 6E/F**), although full processing of caspase-3 leading to PARP cleavage in this model was inhibited (this was expected as HCT116 cells are “Type 2”, meaning they require involvement of the mitochondrial apoptotic pathway to fully activate caspase-3 downstream of the DISC^34^). These effects were maintained at longer timepoints (**Supplementary Figure 7B**), and significantly enhanced TRAIL-induced apoptosis in cells co-treated with MLN4924 was quantitatively confirmed (**Figure 6G**).

### Cullin-1 can regulate the composition and signalling outcomes of the TRAIL-R2 DISC

As MLN4924 affects multiple substrates, including Cullin-3, we assessed to what extent its effects on TRAIL-R2 composition and signalling are Cullin-1-dependent using a Cullin-1-directed siRNA pool (siCul1); the pan-caspase inhibitor (zVAD) was again used to allow us to visualise unprocessed FLIP(L) at the DISC and to inhibit apoptotic cell death induced by prolonged (48h) Cullin-1 silencing, which could have confounding effects on expression of DISC components. Increases in both forms of FLIP(L) and TRAF2 were observed in siCul1-transfected cells, along with a slight increase in FLIP(S) (**Figure 7A**). siCul1 had less impact on poly-ubiquitinated caspase-8 than MLN4924, consistent with this effect of the NEDDylation inhibitor being predominantly mediated via Cullin-3^24^, and its impact on RIPK1 levels was small. Collectively, these results indicate that the interaction of Cullin-1 with the TRAIL-R2 DISC decreases the levels of p43-FLIP and TRAF2 associated with the complex. The impact of these changes on TRAIL-induced apoptosis was similar to those observed for MLN4924, with increased levels of cleaved PARP and p18-caspase-8 (**Figure 7B**) and significantly enhanced cell death induction (**Figure 7C**). In HCT116 cells, siRNA-mediated down-regulation of FLIP(L) inhibited procasapse-8 processing at the TRAIL-R2 DISC thereby reducing the levels of fully processed p18-caspase-8 detected in the unbound fraction (**Figure 7D**). This indicates that FLIP(L) facilitates procaspase-8 processing in this setting and is consistent with the impact of MLN4924 treatment and Cullin-1 silencing, both of which enhance FLIP(L) levels at the TRAIL-R2 DISC and promote TRAIL-induced apoptosis in this model.

**Figure 7.**
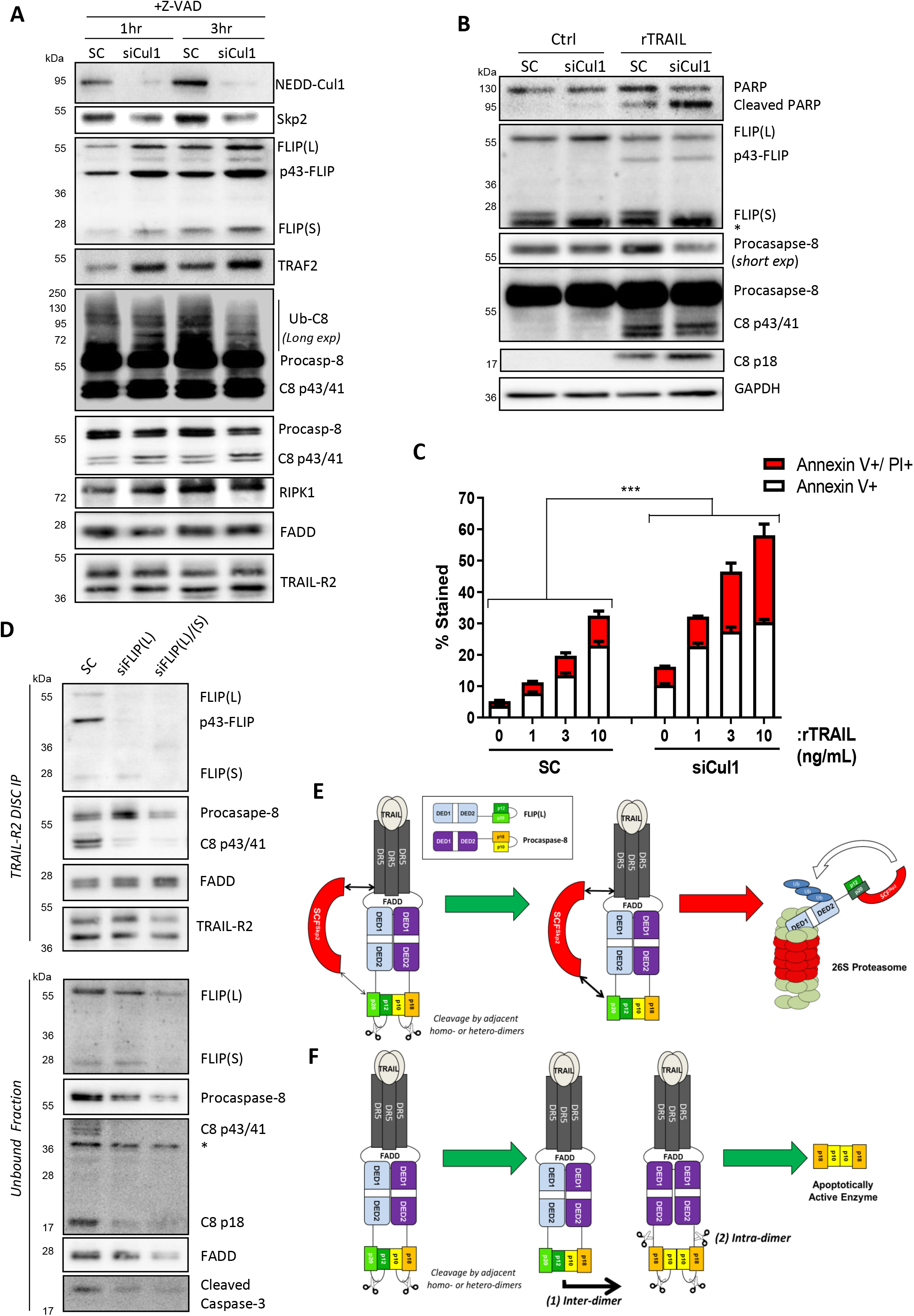
SCF^Skp2^ inhibits TRAIL-induced apoptosis in HCT116 cells. **(A)** Western blot analysis of TRAIL-R2 DISC IP in HCT116 cells transfected for 48h with 30nM control (SC) or a pool of Cullin-1 targeted (siCul1) siRNAs prior to treatment with z-VAD-fmk (20μM) for 2h and then activation of the TRAIL-R2 receptor for 1 and 3h. **(B)** Western blot analyses of apoptosis proteins in HCT116 cells transfected with 10nM control (SC) or Cullin-1-targeted (siCul1) siRNAs for 48h prior to treatment with rTRAIL (10ng/mL) for 6h. **(C)** High content microscopy analysis of Annexin V/PI staining in HCT116 cells transfected with 10nM control (SC) or Cullin-1-targeted (siCul1) siRNAs for 24h prior to treatment with rTRAIL (10ng/mL) for an additional 24h. Significance was calculated by a Two-way ANOVA test **(D)** Western blot analysis of TRAIL-R2 DISC IP in HCT116 cells transfected with 10nM of a pool of FLIP(L)-specific or a pool of pan-FLIP siRNAs and treated with 10μM z-VAD-fmk for 16h prior to DISC stimulation for 1h. **(E)** *Schematic overview of regulation of FLIP(L) by SCF^Skp2^ at the TRAIL-R2 DISC:* cleavage of FLIP(L) to p43-FLIP at the TRAIL-R2 DISC (by proximal FLIP(L)/caspase-8 heterodimers or caspase-8 homodimers) enhances its interaction with Cullin-1, which disrupts its interaction with caspase-8 and promotes its ubiquitination and proteasomal degradation by SCF^Skp2^. This is facilitated by the constitutive interaction of SCF^Skp2^ with TRAIL-R2 and reduces the levels of the FLIP(L):caspase-8 heterodimeric enzyme at the DISC. **(F)** In the absence of Cullin-1, the FLIP(L):caspase-8 heterodimer promotes inter-dimer cleavage (*1*) between the small and large catalytic domains of proximal dimers. In the case of procaspase-8:procaspase-8 homodimers, this is followed by rapid intra-dimer cleavage (*2*) resulting in full processing of procaspase-8 to the pro-apoptotic p10/p12 hetero-tetramer.

## Discussion

FLIP is a major apoptosis-regulatory protein frequently overexpressed in solid and hematological cancers and is an important mediator of chemo- and radio-resistance as well as apoptosis induced by immune effector cells and therapeutic agonists^35–38^. For these reasons, high FLIP expression has been correlated with poor prognosis in a number of cancers. FLIP is also critical for the survival of Tregs and MDSCs^39,40^, key immunosuppressive cells in the tumor microenvironment. Moreover, through its roles (amongst others) in regulating activation-induced cell death (AICD) during the resolution of acute inflammatory responses, direct regulation of the NLRP3 and AIM2 inflammasomes^41^ and survival of cardiomyocytes^42,43^, FLIP has been linked to a number of other diseases and pathologies; therefore, its precise regulation is vital for normal homeostasis.

The turnover of both FLIP splice forms is regulated by the UPS, with half-lives of ~45 minutes for FLIP(S) and ~3 hours for FLIP(L), which is indicative of distinct regulatory mechanisms for each splice form^6^. Although FLIP(S) lacks the pseudo-caspase C-terminal domain of FLIP(L), both proteins share the K192 and K195 residues that were identified as key ubiquitination sites^6^. A cytoplasmic complex termed the “FADDosome” formed in response to the antimetabolite 5-Fluorouracil was recently described containing FLIP, p53, ATR, caspase-10 and TRAF2, in which TRAF2 ubiquitinates FLIP(L) causing its degradation and activation of caspase-8-dependent apoptosis^44^. ER stress leading to JNK-mediated activation of ITCH was reported to lead to ubiquitination and degradation of FLIP(L)^9,10^. Furthermore, recent studies suggest that the deubiquitinase (DUB) USP8 may interact with ITCH to regulate the relative expression of both splice forms. In this paper, we provide the first evidence for specific control of the long FLIP splice form by the SCF^Skp2^ Cullin-RING E3 Ligase complex.

The Cullin RING Ligase (CRL) family of E3 ligases account for roughly 20% of all proteasomal degradation within the cell^45,46^, and many CRL proteins, especially the F-box family of proteins, have been linked to tumourgenesis^47^. NEDDylation is required for CRLs to be active. Using the NEDDylation inhibitor MLN4924 (pevonedistat), we found that FLIP(L) was a potential CRL substrate. While FLIP downregulation in response to MLN4924 has been reported, this effect was observed using higher concentrations of MLN4924 over longer periods than used in our study^48,49^; and indeed we also observed a downregulation of FLIP(L) after 24hrs of treatment (**Supplementary Figure 7B**). Therefore, MLN4924 has a biphasic effect on FLIP(L), with early upregulation followed by downregulation potentially due to suppression of NFκB, a known regulator of FLIP expression^50^ and a known target of MLN49 2 4^49,51,52^. In this study, we focussed on the more direct early effects of MLN4924 on FLIP(L) and the TRAIL-R2 DISC.

Skp2 is an F-box protein linked to TRAIL-induced apoptosis^18^. This led us to us to investigate the most well characterised CRL, the SCF^Skp2^ complex. A specific interaction between SCF^Skp2^ and FLIP(L) but not FLIP(S) was detected in co-IP experiments. Downregulation of SCF^Skp2^ components with siRNA decreased FLIP(L) ubiquitination, while co-overexpression of Cullin-1 and Skp2 increased FLIP(L) ubiquitination in a NEDDylation-dependent manner. Furthermore, MLN4924 treatment stabilised FLIP(L) but not FLIP(S), leading us to the conclusion that SCF^Skp2^ could act as a selective E3 ligase for FLIP(L) and mediate its degradation via the UPS. Mapping experiments revealed that SCF^Skp2^ interacted with amino acids 255-267 within the p20 pseudo-catalytic domain of FLIP(L), a region which is not present in FLIP(S), nor its structurally-related paralogs caspase-8 and −10. Importantly, a stronger interaction with SCF^Skp2^ was detected with the p43-FLIP form of FLIP(L), which is usually generated by caspase-8 in FADD-dependent complexes such as the DISC. Molecular modelling studies using the crystal structure of the FLIP(L):procaspase-8 heterodimer indicated that the Cullin-1 binding site in FLIP(L)’s p20-subunit is partly buried in the interface with its p12-subunit, suggesting that cleavage between these domains by caspase-8 at the DISC may enhance the surface exposure of the Cullin-1-binding domain.

Notably, we found that SCF^Skp2^ could interact with TRAIL-R2 in a manner not dependent on FLIP or other canonical DISC components, FADD and caspase-8. Moreover, this interaction appears to be with the TRAIL-R2 pre-ligand association complex (PLAC) as it does not require receptor activation. Thus, our data suggest a model in which processing of FLIP(L) to p43-FLIP at the TRAIL-R2 DISC enhances its interaction with co-localised SCF^Skp2^, leading to targeting of p43-FLIP to the proteasome and reduced levels of the FLIP(L):caspase-8 heterodimer (**Figure 7E**). This is consistent with the observed longer half-life of non-cleavable FLIP(L) and our observation that overexpressed Cullin-1 was able to disrupt p43-FLIP’s interactions with FADD and caspase-8, indicative of competition between the SCF^Skp2^ complex and canonical DISC proteins for p43-FLIP binding. In this model in the absence of Cullin-1 (**Figure 7F**), enhanced levels of the FLIP(L):caspase-8 heterodimeric enzyme at the TRAIL-R2 DISC would promote processing of its local substrates, including proximal procaspase-8 homodimers, thereby facilitating full processing of procaspase-8 to its pro-apoptotic p10/p18-form.

Canonically, SCF complexes interact with their substrates through their F-box protein; however, we found that p43-FLIP interacts directly with the CR1 domain of Cullin-1, independently of Skp1 and Skp2. Thus, FLIP(L)’s interaction with SCF^Skp2^ is more similar to other proteins known to regulate CRLs, such as CAND1 and subunits of the CSN complex (COP9 signalosome). SCF complexes are capable of auto-ubiquitination (a characteristic shared by many E3 ligases) and can degrade Skp2 in this manner^53^; this is thought to be a negative feedback mechanism that stops these E3 ligases becoming overactive. Similarly, the direct interaction of FLIP(L) with Cullin-1 may trigger its ubiquitination. Direct binding of FLIP(L) to Cullin-1 may also disrupt Cullin-1’s interactions with other substrates and may explain why FLIP(L) overexpression has been observed to inhibit β-catenin proteasomal degradation, which is induced by the Cullin-1-containing SCF^β-TrCP^ complex^54,55^.

Of note, Cullin-1 was predominantly detected in its NEDDylated (active) form at the TRAIL-R2 DISC, where we also detected the previously reported Cullin-3^27^. Inhibition of NEDDylated Cullin-1 using MLN4924 or downregulation of total Cullin-1 with siRNA was found to increase p43-FLIP levels at the DISC. By forming an enzymatically active heterodimer with p43/41-caspase-8 at the DISC, p43-FLIP can inhibit apoptosis when present at high levels or promote apoptosis when present at low levels. In HCT116 cells, we found that siRNA-mediated downregulation of FLIP(L) inhibited procaspase-8 processing at the TRAIL-R2 DISC indicative of a pro-apoptotic role for FLIP(L) at the level of DISC stimulation used in these studies. Consistent with this, downregulation of NEDDylated- and total Cullin-1 using MLN4924 and siRNA respectively enhanced TRAIL-induced apoptosis in these cells, suggesting that by stabilizing p43-FLIP at the DISC, loss of Cullin-1 promotes p43-FLIP-mediated processing of procaspase-8 at the DISC and apoptosis induction. Moreover, the FLIP(L):caspase-8 heterodimer inhibits necroptosis by cleaving RIPK1, and we found reduced levels of RIPK1 at the DISC in the presence of MLN4924 (although not siCul1), suggestive of enhanced cleavage.

FLIP(L)’s TRAF2 binding site, which we mapped to the linker region between DED2 and its p20 pseudo-caspase subunit, is adjacent to the Cullin-1 binding site, and although both can bind simultaneously (**Figure 4D**), the binding of TRAF2 is clearly weakened by Cullin-1 (**Figure 4E**), suggesting competition between the 2 proteins for binding to DISC-processed FLIP(L). Thus, in cells treated with MLN4924 or in which Cullin-1 was silenced, DISC recruitment of TRAF2 was enhanced along with FLIP(L). Our findings (**Figures 6B/C**) agree with those of others^56,57^ that TRAF2 forms part of a FADD- and caspase-8-independent TRAIL-R2 PLAC and suggest that like other E3 ligases, c-Cbl/Cbl-b and A20, SCF^Skp2^ can also associate with the TRAIL-R2 PLAC. Moreover, our findings agree with earlier studies which reported that TRAF2 interacts with p43-FLIP at the DISC, leading to the latter’s stabilisation^58^.

In conclusion, we have identified SCF^Skp2^ as a key regulator of FLIP(L) ubiquitination and stability. Moreover, FLIP(L) can interact directly with Cullin-1; hitherto, the only direct Cullin-1-interacting proteins identified were Skp1, Rbx1, CAND1 and CSN2 and CSN4 of the COP9 Signalosome. Moreover, we have found that SCF^Skp2^ can interact with both the TRAIL-R2 PLAC and DISC. Whereas Cullin-3 has been reported to regulate the TRAIL-R2 DISC by modulating caspase-8, our results indicate that Cullin-1 regulates the DISC by modulating FLIP(L). Thus, our findings add important layers to our understanding of how FLIP expression and TRAIL-R2 signaling are controlled with implications for therapeutically modulating cell death

## Supplementary Figure Legends

**Supplementary Figure 1.**
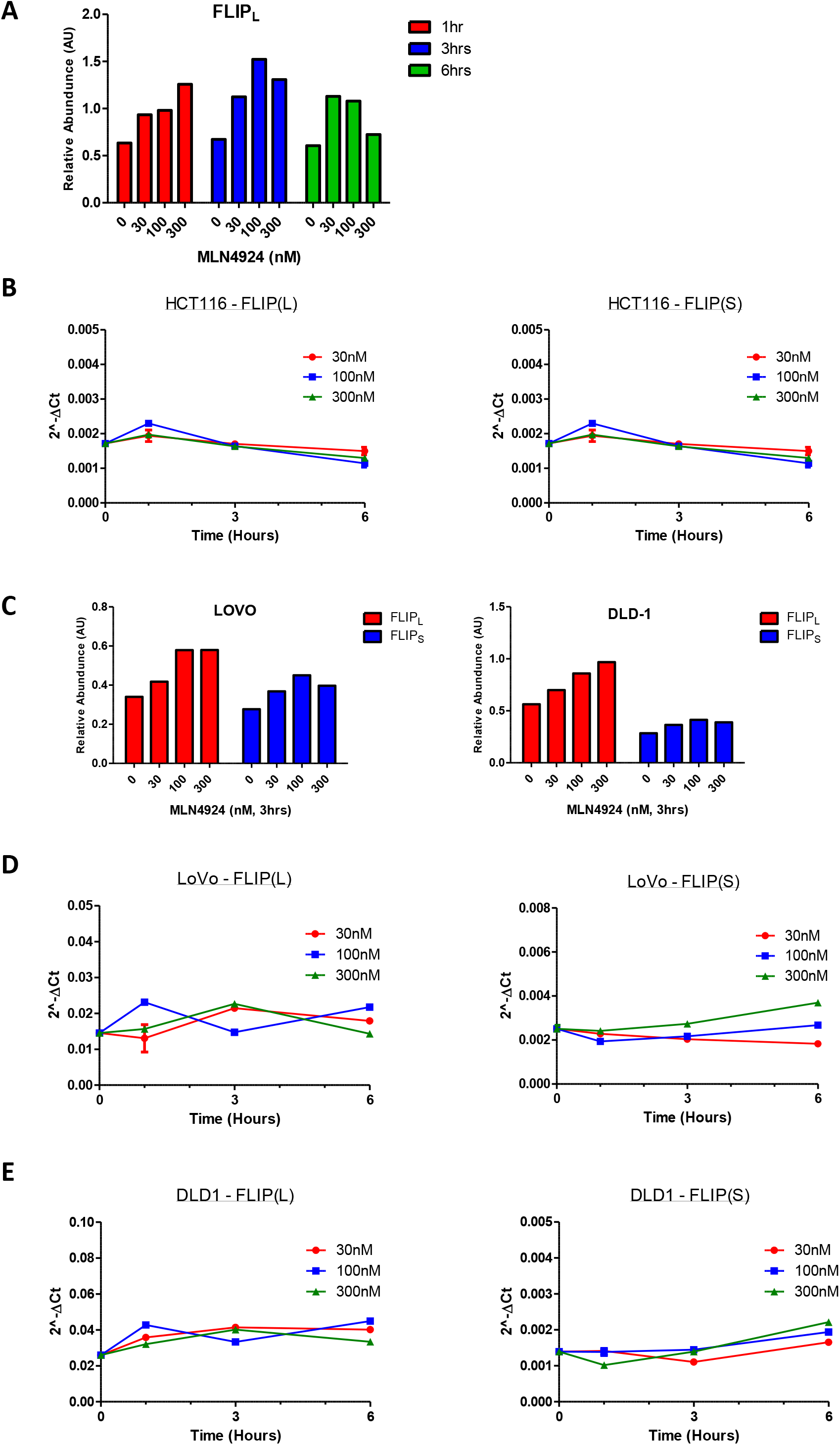
Impact of MLN4924 on FLIP. **(A)** Densitometric quantification of FLIP(L) in the Western blots shown in *Figure 1B*. **(B)** PCR analyses of FLIP mRNA expression in HCT116 cells treated with MLN4924. **(C)** Densitometric quantification of FLIP(L) in the Western blots shown in *Figure 1C*. PCR analyses of FLIP mRNA expression in **(D)** LoVo and **(E)** DLD1 cells treated with MLN4924.

**Supplementary Figure 2.**
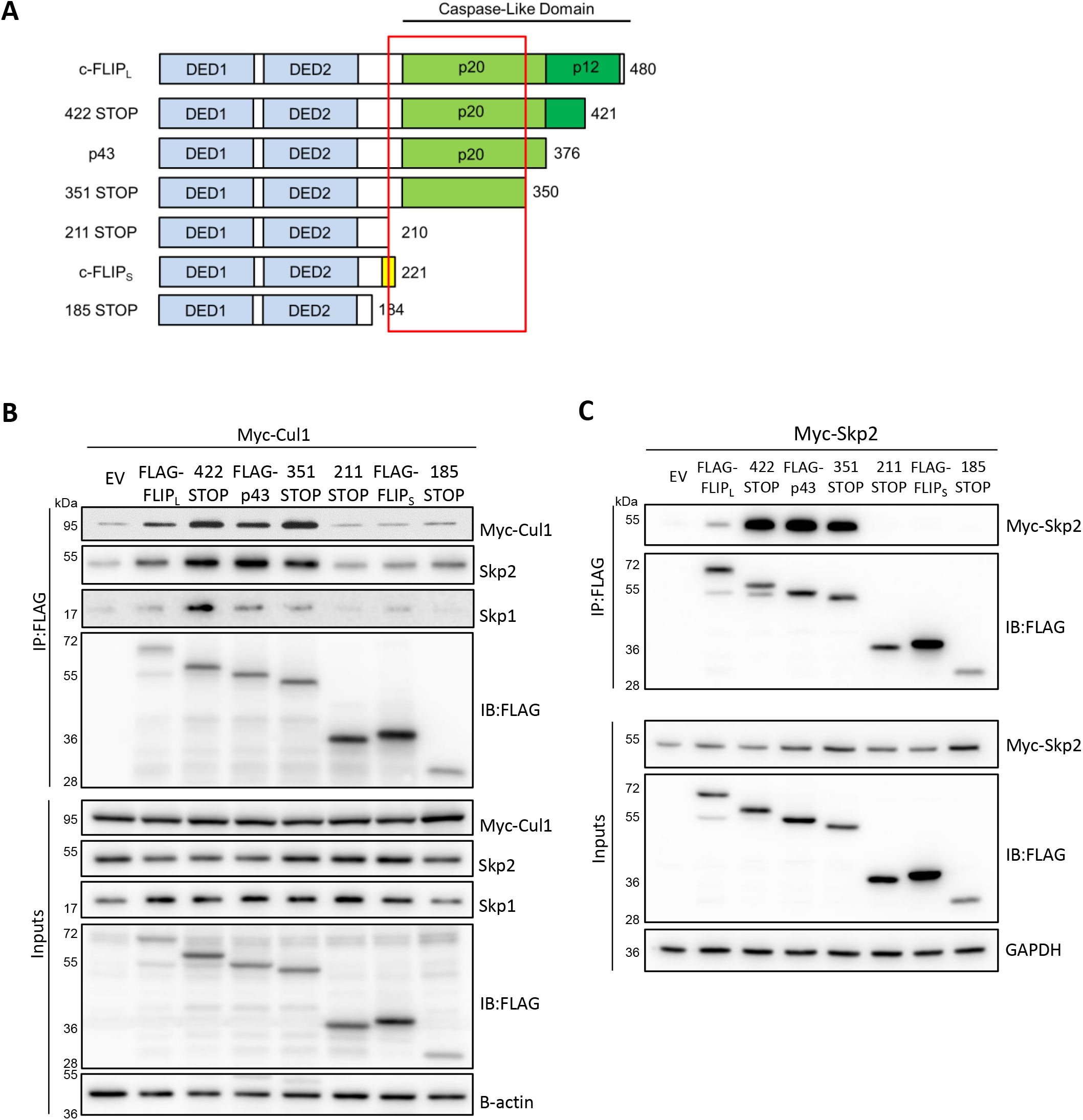
Mapping the interaction between FLIP(L) and the SCF^Skp2^ complex. **(A)** Schematic diagram of FLIP expression constructs; the death effector domains (DEDs) and large (p20) and small (p12) subunits of the pseudo-caspase domain are highlighted. Co-IP experiment mapping the interaction site between FLIP(L) and **(B)** Cullin-1 and **(C)** Skp2.

**Supplementary Figure 3.**
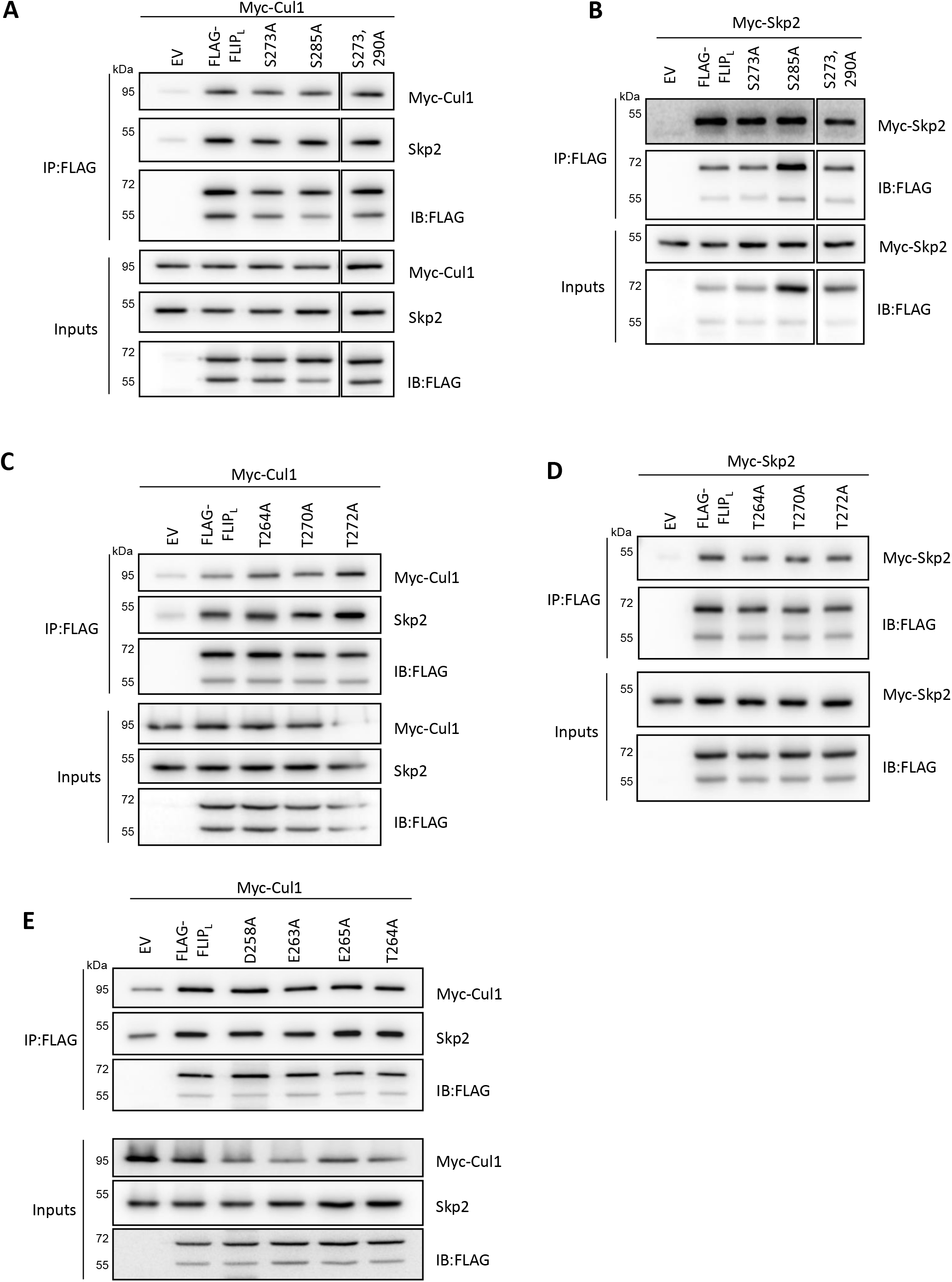
Fine mapping of the interaction between FLIP(L) and the SCF^Skp2^ complex. **(A-E)** Co-IP experiments assessing the importance of candidate serine, threonine, glutamic acid and aspartic acid residues of FLIP(L) for the interaction site between FLIP(L) and Cullin-1 and Skp2.

**Supplementary Figure 4.**
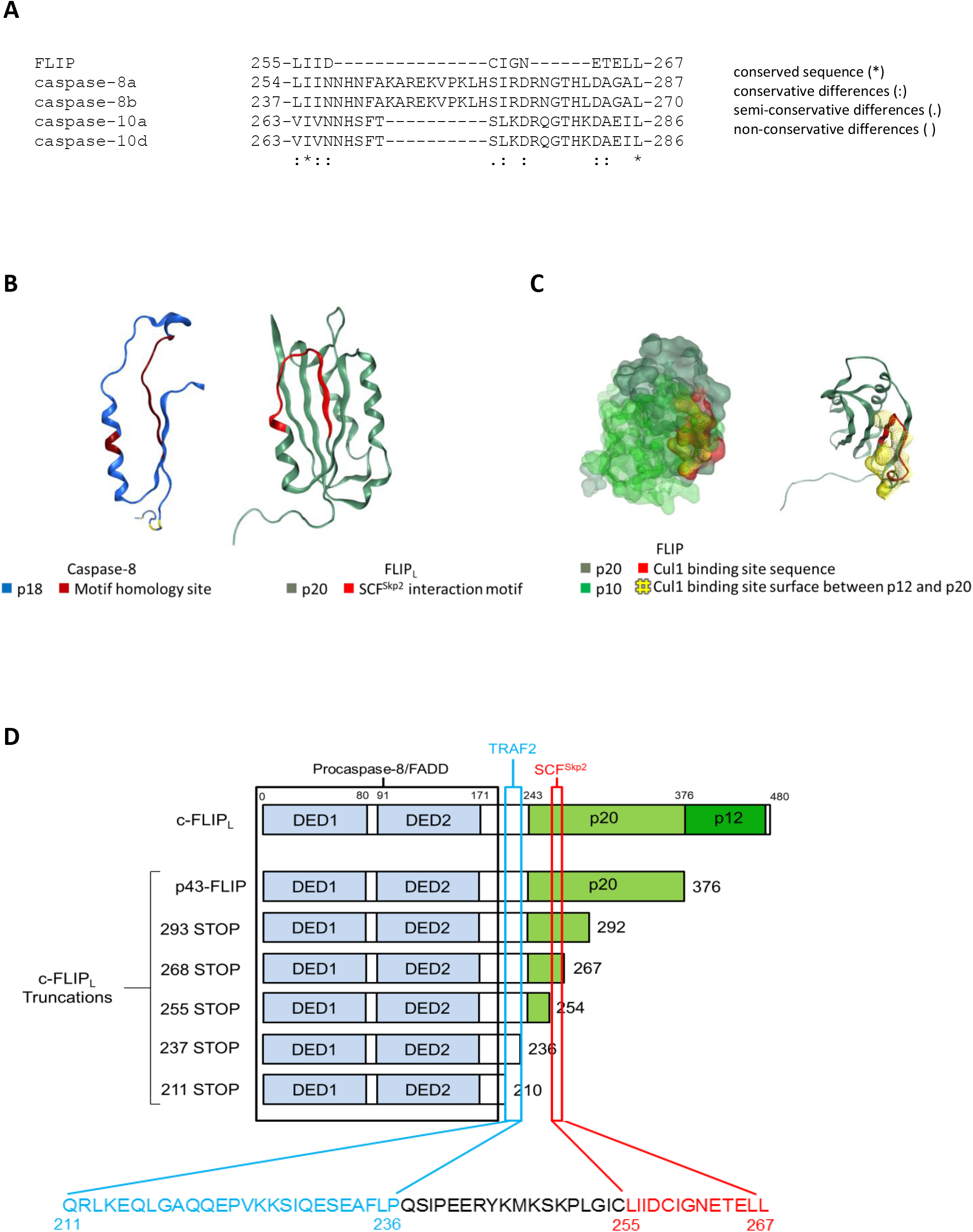
Analysis of the interaction site of FLIP(L) with the SCF^Skp2^ complex. **(A)** Alignment of the Cullin-1 interaction site in FLIP(L) with the corresponding regions in its paralogs caspase-8 and caspase-10. **(B)** Ribbon models of the Cullin-1 interaction site in FLIP(L) and the corresponding region of caspase-8. **(C)** Space-filling and ribbon models of the Cullin-1 interaction site in the large p20 subunit of FLIP(L) and its spatial orientation with respect to the p12 small subunit. **(D)** Schematic diagram of FLIP(L) highlighting its sites of interaction with the DISC components procaspase-8 and FADD and with TRAF2 and SCF^Skp2^.

**Supplementary Figure 5.**
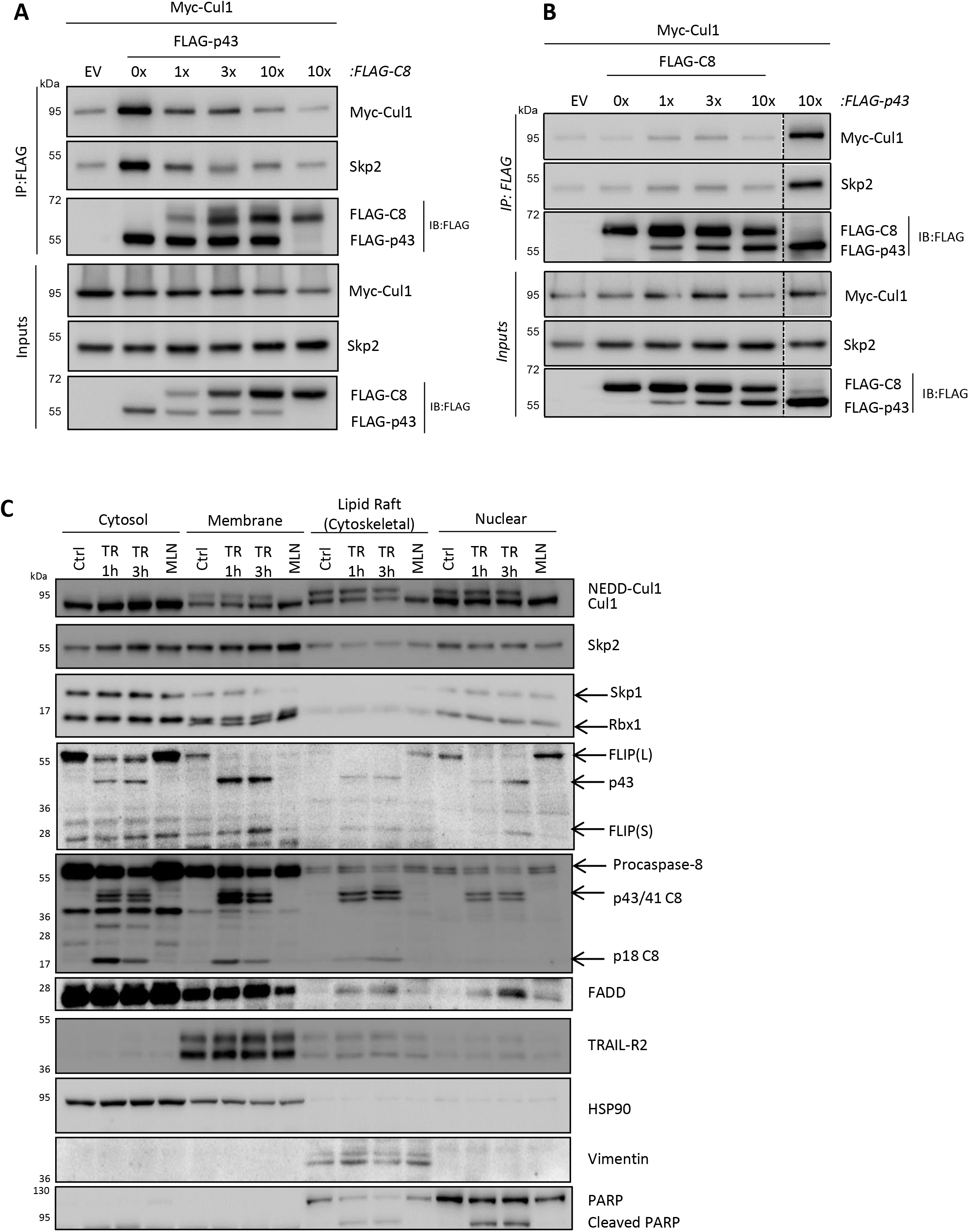
Competition between SCF^Skp2^ and procaspase-8 for p43-FLIP binding and subcellular fractionation studies. **(A)** Co-IP analysis of the interaction of Myc-tagged Cullin-1 and endogenous Skp2 with Flag-tagged p43-FLIP in cells co-transfected with increasing amounts of Flag-tagged procaspase-8. **(B)** Co-IP analysis of the interaction of Myc-tagged Cullin-1 and endogenous Skp2 with increasing amounts of Flag-tagged p43-FLIP in cells co-transfected with Flag-tagged procaspase-8. **(C)** Western blot analysis of sub-cellular expression of SCF^Skp2^ and TRAIL-R2 DISC proteins in HCT116 cells treated with 10ng/mL IZ-TRAIL for the indicated times or 1μM MLN4924 for 3h.

**Supplementary Figure 6.**
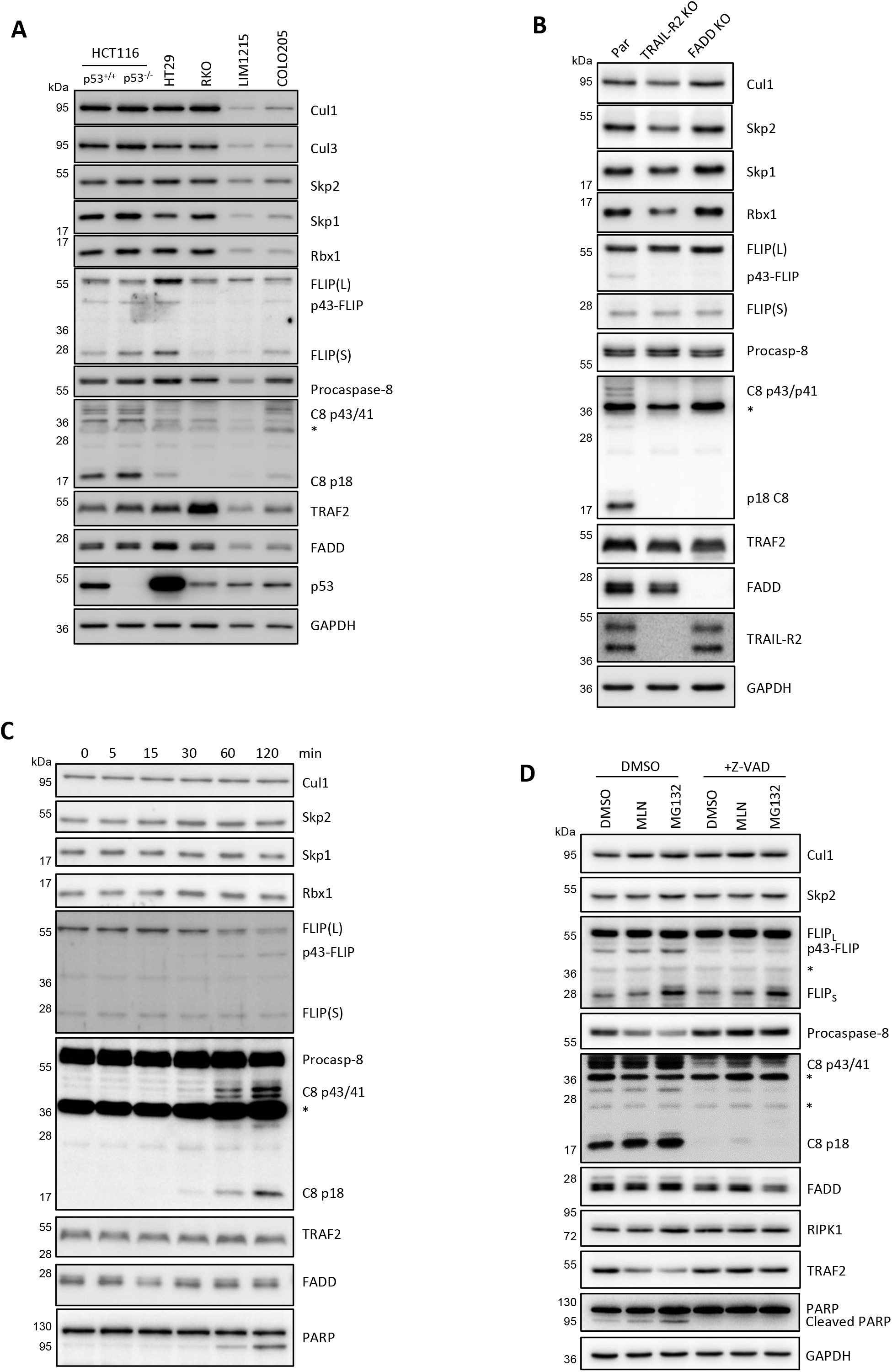
Unbound fractions from the DISC IPs in *Figure 6*.

**Supplementary Figure 7.**
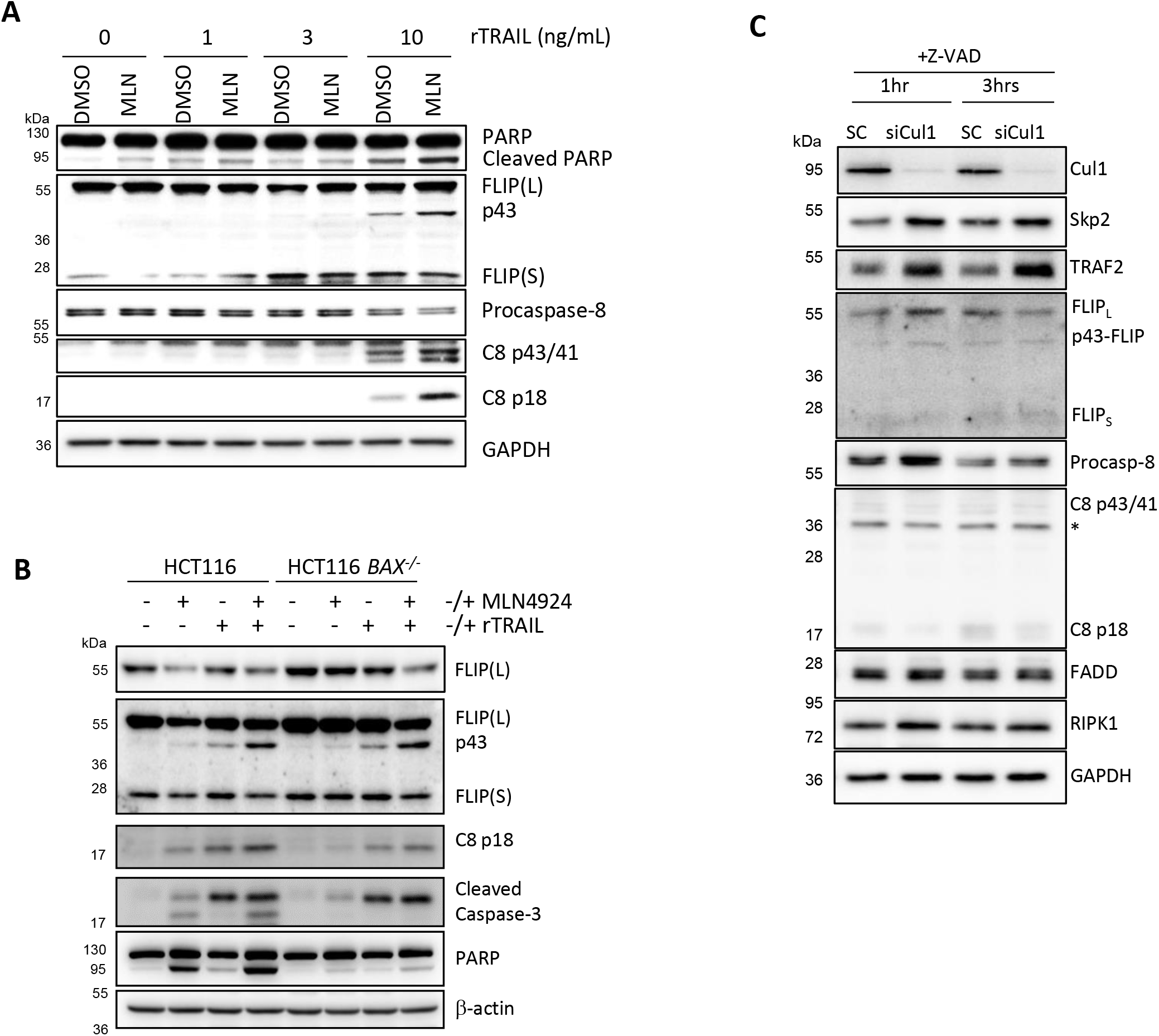
SCF^Skp2^ inhibits TRAIL-induced apoptosis in HCT116 cells. **(A)** Western blot analyses of apoptosis proteins in HCT116 parental cells pre-treated with MLN4924 (100nM) for 1h prior to treatment with rTRAIL for 6h. **(B)** Western blot analyses of apoptosis proteins in HCT116 parental and Bax-deficient cells co-treated with MLN4924 (100nM) and rTRAIL (10ng/mL) for 24h. **(C)** Unbound fractions from the DISC IP in *Figure 7A*.

## Methods

### Cell Culture

HCT116 cells were cultured in McCoy’s 5A Modified Medium, supplemented with 10% fetal calf serum (FCS) and 2mM L-glutamine (all from Life Technologies). HEK293T cells were maintained in Dulbecco’s Modified Eagle Medium (DMEM), supplemented with 10% foetal calf serum (FCS) and 1mM Sodium Pyruvate (all from Life Technologies). DLD-1 and COLO205 cells were maintained in Roswell Park Memorial Institute (RPMI) 1640 Medium (Sigma Aldrich, Gillingham, Dorset, UK), supplemented with 10% FCS (Life Technologies). LOVO, RKO, HT29 and LIM1215 cells were maintained in Dulbecco’s Modified Eagle Medium (DMEM) supplemented with 10% FCS (all from Life Technologies). Cells were maintained in a tissue culture incubator (Sanyo Europe, Herts, UK) at 5% CO_2_ and 37°C.

### Oligonucleotide Transfections

A 3:1 ratio of FuGENE^®^ HD Transfection Reagent (Promega) (μL) to plasmid DNA (μg) was used for transfections. In 100μL of OptiMEM (Life Technologies), FuGENE then plasmid DNA was added and left to incubate for 15 mins at room temperature. The mixture was then added to cells in culture and was left for 24-48h. For siRNA transfections, siRNA and Lipofectamine^®^ RNAiMAX (Invitrogen) were added separately to OptiMEM (Life Technologies) before being mixed together and left to incubate at room temperature for 15 mins. Culture medium was then added to the mixture, to allow total cell coverage, and was used to replace the medium on cells, which was then left to incubate at 5% CO2, 37°C for 4h. Further culture medium was then added, to normal culturing levels, and then cells were left for 16-72h.

### Immunoblotting

Cell lysates were prepared or immunoprecipitation (IP) samples were eluted using Loading buffer, subsequently followed by heating at 95°C for 5 mins. SDS-PAGE was then implemented to separate proteins by molecular weight and were then transferred to a nitrocellulose membrane. Protein expression was analysed by immunoblotting and detected using the G:BOX Chemi XX6 gel doc system (Syngene).

### Ubiquitination Assay

Cells grown in a P90 plate, typically collected at over 80% confluency, were washed once in ice-cold PBS to stop dynamic ubiquitination events from occurring; all procedures were then conducted at 4°C for the same reason. Cells were transferred to tubes and were pelleted by centrifugation at 2400 rpm for 5 mins; supernatant was subsequently removed. Cells were then lysed with 250μL of SDS-free RIPA buffer (50mM Tris pH 7.4, 150mM NaCl, 5mM EDTA, 1% Triton X-100, dH_2_O), supplemented with an EDTA free protease inhibitor cocktail, for 20 mins before centrifugation at 13,00 rpm for 5 mins, followed by the supernatant being transferred to fresh tubes. 25μL of 10% SDS was added (Final concentration 1%) and the sample was heated at 95°C for 5 mins; to facilitate denaturation of the samples. The samples were then diluted with 750μL of SDS-free RIPA buffer, supplemented with an EDTA free protease inhibitor cocktail, to dilute the SDS and allow the immunoprecipitation (IP) antibody to function efficiently; samples were then incubated with 1μL of either FLIP (H-202) antibody (Santa Cruz biotechnology) or FLAG (M2) antibody (Sigma) overnight at 4°C while rotating. 30μL of the appropriate washed Dynabeads^™^ M-280 (Invitrogen) were then added to the samples and left to incubate for 6h at 4°C while rotating. The beads were then washed 5 times in SDS-free RIPA buffer and eluted in Loading buffer before being heated at 95°C for 5 mins.

### Co-Immunoprecipitation (Co-IP)

Cells grown in a P90 plate, typically collected at over 80% confluency, were washed once in ice-cold PBS to stop disruption of protein:protein interactions; all procedures were then conducted at 4°C for the same reason. 1mL of SDS-free RIPA buffer (50mM Tris pH 7.4, 150mM NaCl, 5mM EDTA, 1% Triton X-100, dH_2_O), supplemented with an EDTA free protease inhibitor cocktail, was added onto cells within the plate and was left shaking for 20 mins. Cell lysates was then transferred to a tube, centrifuged at 13,000 rpm for 5 mins and the supernatant was transferred to a fresh tube. Either 1μL of FLIP (H-202) antibody (Santa Cruz biotechnology)/Myc-Tag (9B11) antibody (Cell Signaling Technology) or 20μL of FLAG (M2) Magnetic Beads (Sigma) were added to the cell lysates and were left to rotate overnight at 4°C. The following day 15μL of the appropriate washed Dynabeads^™^ M-280 (Invitrogen) were added to the samples and were left to rotate for a further 6h. The beads were then washed 5 times with SDS-Free RIPA buffer and were eluted in Loading buffer, followed by heating at 95°C for 5 mins. However, if the FLAG (M2) Magnetic Beads were used the Dynabeads step was skipped and beads were washed straight away.

### TRAIL-R2 DISC IP

1.2mg of anti-TRAIL-R2/DR5 antibody (Conatumumab, AMG655) (Amgen Inc.) was covalently attached to 60mg of Dynabeads in a volume of 6mL using the Dynabead^®^ Antibody Coupling Kit (Life Technologies), according to the manufactory’s instructions. Typically, a P90 plate containing 10mL of growth medium was inoculated with 30μL of anti-DR5 beads when cells were approximately 60% confluent. The cells were then washed once in ice-cold PBS to slow down DISC dynamics and further procedures were carried out at 4°C for the same reason. 1mL of DISC IP buffer (20mM Tris 7.4, 150mM NaCl, 0.2% NP-40, 10% Glycerol, dH_2_O), supplemented with an EDTA free protease inhibitor cocktail, was added onto cells within the plate and allowed to shake for 30 mins, then was transferred to a fresh tube. Anti-DR5 beads were isolated magnetically and washed 5 times with DISC IP buffer; the first flow-through was collected and termed the ‘Unbound Fraction’. Beads were then eluted using Loading buffer and analysed by Immunoblotting. AMG655 is an agonist antibody for TRAIL-R2DR5, therefore, activating the receptor, when it is immunoprecipitated after being added to live cells any complexes that associate with active DR5 (like the DISC) are also immunoprecipitated as well.

### Biotin Pulldown

For each sample, 25μL of Dynabeads^™^ MyOne^™^ Streptavidin T1 (Invitrogen) were first washed in SDS-free RIPA buffer (50mM Tris pH 7.4, 150mM NaCl, 5mM EDTA, 1% Triton X-100, dH_2_O) and then resuspended in SDS-Free RIPA+2% BSA for blocking. 20μg of the Biotin tagged peptide (peptides&elephants) was then conjuagted to the beads at room temperature of 30 mins while rotating. The conjugated beads were then washed with SDS-Free RIPA buffer and incubated with either rCullin-1 (abcam) or 50μg of cell lysate, collected in SDS-Free RIPA buffer supplemented with an EDTA free protease inhibitor cocktail, for 4h at 4°C while rotating. Beads were then washed with SDS-Free RIPA buffer and eluted in Loading buffer, followed by heating at 95°C for 5 mins.

### PCR

A PCR reaction mixture of 3μL of cDNA, 1μL DNase/RNase free water, 0.5μL Forward primer (5μM), 0.5μL Reverse primer (5μM) and 5μL LightCycler^®^ 480 SYBR Green I Master reagent (Roche Diagnostics) was performed and analysed on a Roche LightCycler^®^ 480 System (Roche Diagnostics). Relative mRNA levels were then normalised to the housekeeping gene (RPL24) and determined by the ΔΔCt method.

### Site Directed Mutagenesis

Site directed mutagenesis was carried out using the KOD Xtreme^™^ Hot Start DNA Polymerase kit (Merck). Briefly, a PCR reaction (100μL scale) incorporating 50μL 2x buffer, 20μL of 2mM dNTPs, 6μL each of 5μM forward and reverse primers (containing the desired mutations), 16μL of 10ng/μL template DNA and 2μL KOD polymerase was set up. 50μL of the completed PCR reaction mixture was then digested with 2.5μL Dpn1 and 5.8μL CutSmart^®^ Buffer (New England Biolabs) for 3h at 37°C to degrade methylated template DNA. CaCL_2_ competent DH5a bacteria when then transformed with the PCR mixture and subsequent colonies were picked and verified by DNA sequencing (GATC BIOTECH).

### Caspase Activity Assay

25μL of Caspase-Glo^®^-3/7 or −8 reagent (Promega) was added to 5μg (Caspase-3/7 activity) or 10μg (Caspase-8 activity) of cell lysate, made up to 25μL with PBS in a white-walled 96 well plate in duplicate. The plate was incubated in the dark at room temperature for 45 minutes and subsequently read at 1 sec integrated reads in a luminescent plate reader (Biotek Synergy 4 plate reader).

### High Content Fluorescent Screening

Cells were seeded into a 96-well glass-bottomed plate (Cellvis) and left to adhere. After treatments, cells were incubated with 10x Annexin V Binding Buffer (BD Pharmingen), 1:1000 FITC Annexin V (BD Pharmingen), 0.333μg/mL Propidium Iodine (Sigma Aldrich) and 1.33μg/mL Hoechst 33342 (Thermo Fisher Scientific) for 20 mins at room temperature. The plate was then analysed on the ArrayScan^™^ XTI HCA Reader, integrated with the CrEST^™^ X-Light^™^ Confocal Scan Head (Thermo Fisher Scientific), and the HCS Studio Cell Analysis Software V6.6.0 (Thermo Fisher Scientific) was used to calculate the percentage of cells stained with Annexin V and Propidium Iodine. A target of 2000 cells was read in each well and all conditions were performed in triplicate.

### Sub-Cellular Fractionation

The ProteoExtract^®^ Subcellular Proteome Extraction Kit (Calbiochem) was used to generate lysates from the cytosolic, nuclear and cytoskeletal/lipid raft fractions of the cell, according to the manufacturer’s instructions. The fractions were then subsequently analysed by immunoblotting.

### Protein Half-life

Protein expression was analysed by Immunoblotting and quantified by densitometry, using ImageJ (NIH); normalising to the loading control. Treatment values were converted into ratios of its 0h timepoint (given the value 1) and were plotted. GraphPad Prism 5 was then used to generate a decay curve for treatments, by using the one phase exponential decay equation.

### Statistical Analysis

Statistical significance was calculated from distinct technical replicates (n=3) by Two-way ANOVA with a Bonferroni post hoc test performed in GraphPad Prism 5. Graphs were plotted as means with error bars represented as SEM; statistical significance was denoted as follows: *** = p < 0.001, ** = p < 0.01, * = p < 0.05, ns = p>0.05.

## Acknowledgements

This work was funded by grants from The Wellcome Trust (110371/Z/15/Z), Cancer Research UK (C11884/A24387), Northern Ireland Department for the Economy (NI DfE) (SFI-DEL 14/1A/2582) and a NI DfE studentship (JZR). We thank Dr Jon Vosper (Innsbruck Medical University) for supplying the FLAG-tagged Skp2 expression construct and its corresponding truncation, and Prof Markus Rehm (Stuttgart) for supplying the TRAIL-R2/DR5 knockout HCT116 cell line.

## Author Contributions

JZR: conceptualization, methodology, validation, formal analysis, investigation, writing (original draft) and visualization; CH: conceptualization, methodology, validation, formal analysis, investigation; TS: investigation and visualization; JF: investigation; C Higgins: methodology, investigation and supervision; GE-F: investigation; JM: methodology and investigation; NC: resources and supervision; JSR: investigation; HK: investigation; LMH: investigation; JF: investigation; PM: funding acquisition, writing (review & editing); SSM: funding acquisition, writing (review & editing); DBL: conceptualization, methodology, validation, formal analysis, writing (original draft), visualization, supervision, project administration and funding acquisition.

## Data Availability

The data supporting the findings of the study are available from the corresponding author on reasonable request.

